# Genetically diverse populations hold the keys to climatic adaptation: a lesson from a cosmopolitan raptor

**DOI:** 10.1101/2023.03.17.533108

**Authors:** Hugo Corval, Tristan Cumer, Alexandros Topaloudis, Flavien Collart, Antoine Guisan, Alexandre Roulin, Jérôme Goudet

## Abstract

Although local adaptation influences species distributions, its role in driving evolutionary resilience under climate change remains unclear. Current predictive models focus on genetic adaptation to present climates, providing limited insight into future adaptive capacity. We hypothesise that historical responses to climatic shifts can reveal future adaptive potential. Combining ecological niche modelling and genomic analyses, we investigated spatiotemporal patterns and mechanisms of local adaptation of the Western Palearctic barn owl (Tyto alba). Ecological modelling revealed that barn owls now occupy a broader climatic niche than during the Last Glacial Maximum. Genomic analyses indicated ongoing adaptation, with regions under selection linked to environmental factors across all populations. Our findings demonstrate that local adaptation drives evolutionary changes across populations, enabling colonisation of new habitats and shaping responses to climate change in resident populations. We demonstrate that standing genetic diversity plays a crucial role in adaptation to past, present, and future environmental shifts.

## Introduction

Climatic variations affect biodiversity by impacting individual’s fitness, by driving population differentiation, and ultimately by shaping species distributions^1–3^. The extent of these impacts, however, depends on the speed and intensity of climatic variations^4^: sudden or extreme shifts can lead to local or global extinction if individuals fail to survive or reproduce^5^. In case of more gradual changes, the persistence of populations and species will depend on individuals’ ability to migrate or cope with their new environment^1,6,7^. Through migration, individuals track their suitable conditions in time and space to survive, inducing a shift in species distribution^1^. Individuals can also change physiologically, morphologically, or phenologically to cope with the new local conditions^7^. These changes can occur via phenotypic plasticity, the ability of individual genotypes to produce different phenotypes when exposed to various environmental conditions^8^ or through genetic adaptation, favouring different genotypes better adapted to the local ecological conditions^9^.

Considering the critical role of local adaptation in determining population persistence^10^, understanding whether individuals possess the intrinsic capacity to adapt to new environmental conditions is critical. In recent years, genomic offset has emerged as a key metric for predicting genetic maladaptation to future climates by linking environmental factors to allele frequencies and estimating the genetic changes needed for individuals to survive to new conditions^11–13^. However, while such predictions provide valuable insights into potential risks, these predictions assume populations are adapted to current conditions and cannot evolve further, overlooking their adaptive potential. Therefore, a key challenge remains understanding whether standing genetic variation can fuel adaptation to new climate - a question our study seeks to answer.

At the end of the Last Glacial Maximum (LGM) temperatures rose and ice caps melted, allowing species to (re-)colonise previously unsuitable lands^2,14,15^. Climatic variations since the LGM offer an excellent opportunity to study species’ adaptation. The interplay between migration and selection gives rise to several possible scenarios for how local adaptation may occur: In populations that migrated northward, individuals either tracked their suitable niche or faced new conditions. In the latter case, selection may have driven local adaptation by favouring the most suited individuals^9^, but repeated migration events eventually caused founder effects and a loss of adaptive potential along the front of colonisation^16^. In contrast, populations that remained at the core of the distribution may also have faced a change in conditions. These populations often harbour a higher genetic diversity^17^, possibly enhancing their ability to adapt to their changing environment. Given these three main scenarios, where and how local adaptation happens remains elusive and exploring these questions can enhance our understanding of the adaptive potential of populations.

The barn owl (Tyto alba), a nonmigratory raptor distributed all over the Western Palearctic, faces heterogeneous climates. The species recolonised the Northern part of Europe at the end of the LGM from two main glacial refugia located in the Iberian Peninsula and the Italian and Balkan Peninsulas^15^. Presently, its range stretches across the Western Palearctic^15^. Despite a generally low level of genetic differentiation across its range, southern populations host a higher genetic diversity than northern populations^15,18,19^, and exhibit notable phenotypic variation, such as a cline in plumage coloration between southern and northern populations^18–21^. A previous study demonstrated that the colour cline cannot be explained by purely neutral processes and argued that selection for local adaptation was or is still acting on this phenotype^21^. The well-understood history of the barn owl in Europe since the Last Glacial Maximum, combined with the low genetic structure at the continental level and previous evidence of adaptive selection throughout its range, make this species an attractive model to test and quantify the extent and location of local adaptation.

Here, we evaluated where and how heterogeneous climate induce local adaptation using the European barn owl as a model organism. We first used ecological information and Species Distribution Modelling to (i) quantify the climatic heterogeneity faced by the barn owl nowadays and (ii) measure how these climatic conditions differ from those experienced during the Last Glacial Maximum. We then looked for an association between climatic variables and genomic variants from the entire genome of 74 owls from 9 different localities across Europe. We combined these results with a new approach to scan the genomes for traces of selection and identified genomic regions and genes potentially involved in the local adaptation of the different populations. We found a strong and common signal in southern populations. Overall, our results demonstrate how the most diverse populations, often located at the core of the distribution, may host the adaptive potential to face climate change, giving clues on how standing genetic variation can fuel local adaptation.

## Results

### Suitable conditions nowadays are more diverse than during the Last Glacial Maximum

The spatial prediction obtained by the species distribution model (SDM)^22^ highlights a striking increase of the area occupied by the barn owl since the Last Glacial Maximum^23^ (LGM, ∼ 20,000 years ago), similar to what was observed in trees^24^, mammals^25^, and birds^26^. During the LGM, suitable habitats were mostly confined to regions around the Mediterranean — covering northern Africa as well as the Iberian, Italian, and Greek peninsulas (Supplementary Figure 1). Today, favourable conditions extend well into Western, Central, and Northern Europe, making most of the continent suitable for barn owls. This notable northward and inland expansion prompted us to investigate the climatic variables driving these changes.

To identify the climatic factors defining the climatic niche of the barn owl, we performed a Principal Component Analysis (PCA) fitting a multidimensional climatic space both from the LGM and modern periods. The first three principal components explain 91.50% of the variance (Figure 1b). The first component, explaining 49.66% of the variance, is driven by temperature-related variables - such as the Mean Temperature of the Coldest Month (bio6), Temperature Annual Range (bio7), and Mean Temperature of the Wettest Quarter (bio8) - which differentiate the Mediterranean region from the rest of Europe (Figure 1c). The second component, explaining 33.6% variance, is mainly influenced by precipitation factors, including Precipitation Seasonality (bio15), Precipitation of the Driest Quarter (bio17), and Precipitation of the Coldest Quarter (bio19), effectively distinguishing coastal areas from the continental interior (Figure 1c). The third component, explaining 8.25% variance, is largely driven by the Minimum Temperature of the Coldest Month (bio6; Supplementary Figure 2 and 3).

**Figure 1.**
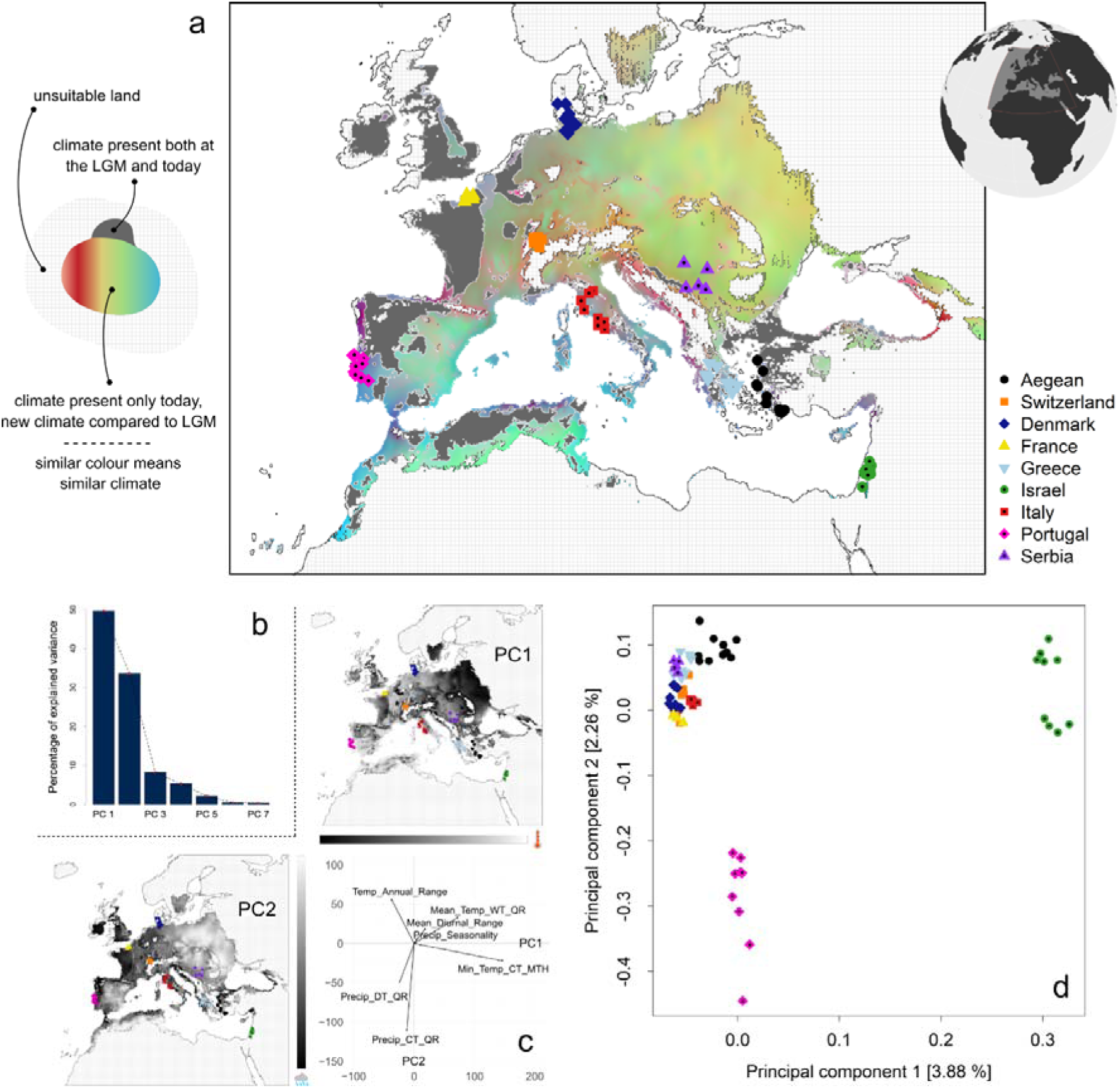
Environmental and genetic variation across the suitable range of the European barn owl. (**a**) Map depicting the climatic heterogeneity across the range of the European barn owl according to the species distribution modelling. Dark grey cells surrounded by a white border have suitable climatic conditions similar to conditions present during the Last glacial Maximum (LGM). Coloured cells outside the white-bordered polygon have climatic conditions not present during the LGM. Colours are based on the multidimensional climatic space of bioclimatic data shown in (b): the scores of the first three principal components (PCs) were converted into values of RGB (PC1: red; PC2: green; PC3: blue) to represent variation in climate. Similar colours represent similar climates. Symbols represent sampling coordinates of individuals from 9 different populations. A jitter has been added for better visualisation (longitude: 0.425; latitude: 0.42). (**b**) Variance explained by the 7 first principal components of the PCA made on bioclimatic variables from the entire study area pictured on (a). Climatic variables came both from the LGM and today to picture the overall climatic variability. (**c**) Correlation between climatic variables and the two first axes of the PCA. Dark to white gradients picture the contribution of each axis projected at the European scale - Abbreviations : CT = Coldest; DT = Driest; MTH = Month; Precip = Precipitation; QR = Quarter; Temp = Temperature; WT = Wettest (d) PCA based on the whole genome of the 74 European barn owls identified in (a). Symbols legend is the same as for panel (a). Only the two first principal components are represented.

We next explored the overlap between past and present climatic conditions in areas suitable for barn owls and showed that owls now occupy a wider variety of climatic conditions than during the LGM^24^. We found that only a small fraction of today’s suitable habitats shares characteristics with those from the LGM. We found that only 25.54% of modern suitable cells exhibit LGM-like climatic conditions (Figure 1a, dark grey areas). These overlapping regions are found in northern Africa, the northwestern Iberian Peninsula, western France, the British Isles, and western Turkey. In contrast, 74.45% of current suitable cells are characterized by climatic conditions that did not exist during the LGM (Figure 1a, RGB-coloured areas). This significant shift suggests that most of today’s barn owl habitats are defined by new climatic conditions, potentially driving local adaptation to these emerging environments.

### Species-wide sampling reveals that a substantial portion of the genome is under climatic selection

To test the hypothesis that local climates may have driven the local adaptation of the barn owl in the Western Palearctic, we have sampled and sequenced 79 individuals from nine populations across the climatic and European geographical distribution of the species (Supplementary Figure 4 and 5, Supplementary Table 1) and identified 12,309,943 Single Nucleotide Polymorphisms (SNPs). The overall differentiation was low (overall F_ST_ = 0.034) and in line with previous estimates^15^. The first axis of the genomic PCA (explaining 3.88% of the total variance) contrasted individuals from the Levant populations (Israel—IS) to all other individuals (Figure 1d). The second axis of the PCA (explaining 2.26% of the variance) opposed individuals from the most diverse population (from the Iberic peninsula—PT) to all others (Figure 1d, Table 1). Overall, southern populations (PT, IT, GR, AE, IS) were more genetically diverse than northern populations (CH, FR, DK, SB) (Table 1). Next, we employed two complementary approaches - a genome scan for selection and a Genotype-Environment Association (GEA) analysis - to pinpoint genomic regions potentially influenced by these climatic conditions.

**Table 1.** Genetic diversity estimated for 9 populations of Western Palearctic barn owls. Standard deviations are found between brackets. See Material and Methods for details on calculation. N, sample size; # PolymorphicS, number of polymorphic sites; # PrivateA, number of private alleles; # RareA, number of rare alleles; PopSpecificFST, population-specific FST computed at the whole genome level.

| Population | N | # PolymorphicS | # PrivateA | # RareA | PopSpecificFST |
| --- | --- | --- | --- | --- | --- |
| AE | 10 | 4,231,387 (44,618) | 193,178 (16,440) | 687,423 (28,433) | 0.03 |
| CH | 9 | 4,259,867 (19,877) | 170,886 (4,123) | 642,071 (9,990) | 0.038 |
| DK | 10 | 4,102,932 (17,633) | 135,566 (3,939) | 560,136 (9,451) | 0.049 |
| FR | 4 | 3,722,772 (0) | 124,430 (1,292) | 487,175 (1,859) | 0.045 |
| GR | 9 | 4,176,443 (19,842) | 144,856 (3,756) | 606,164 (8,453) | 0.046 |
| IS | 9 | 4,402,055 (12,195) | 626,578 (7,595) | 1,173,410 (6,991) | 0.009 |
| IT | 9 | 4,120,120 (10,620) | 214,759 (2,352) | 679,772 (5,274) | 0.054 |
| PT | 9 | 4,678,275 (25,247) | 492,983 (7,476) | 1,090,608 (12,569) | -0.018 |
| SB | 5 | 4,005,667 (0) | 106,496 (2,004) | 514,405 (2,428) | 0.06 |

In our first approach, we scanned the genomes for signatures of selection using population-specific FST^27^. This metric, which accounts for population structure, enables the detection of regions of the genome where individuals have higher genetic similarity than what is observed along the rest of the genome - a pattern possibly resulting from selective events^27,28^. Out of 52,429 windows of 100 kbp (with a 20 kbp slide), we identified 10,607 outlier windows: 7,034 were unique to a single population, and 3,573 were shared between at least two populations. On average, each population had about 1,756 outlier windows (Table S2), with Israel and France at the extremes (989 and 2,087 outliers, respectively).

To ensure that selection signals were caused by climatic conditions, we directly linked genetic variation to temperature and precipitation conditions with a Genotype-Environment Association (GEA). We conducted a Redundancy Analysis (RDA, Supplementary Figure 6)^29^ that we integrated into genomic windows with the Weighted-Z Analysis (WZA) method^30^. We detected 2,181 outlier windows significantly linked to temperature and precipitation variables (Supplementary Figure 7) and overlaid it with the population-specific FST scans.

This combined analysis yielded a refined list of 1,246 outlier windows (displayed in dark blue in Figure 2a; Table S2). 59.71% were unique to a single population while the rest was shared between at least two populations. By merging successive outlier windows, we delineated 270 genomic regions, many forming distinct adaptive peaks along the genome. Particularly striking associations were observed on Super-Scaffold 14 and 45 (Figure 2b). Additionally, we detected a high density of outlier windows in the first half of Super-Scaffold 22 (Figure 2b-d); this signal, shared by owls from France, Italy, and Portugal, corresponded with a marked increase of population-specific FST in these genomic regions (Supplementary Figure 8), indicating that individuals within each of these populations are significantly more similar to one another than to the rest of the samples.

**Figure 2.**
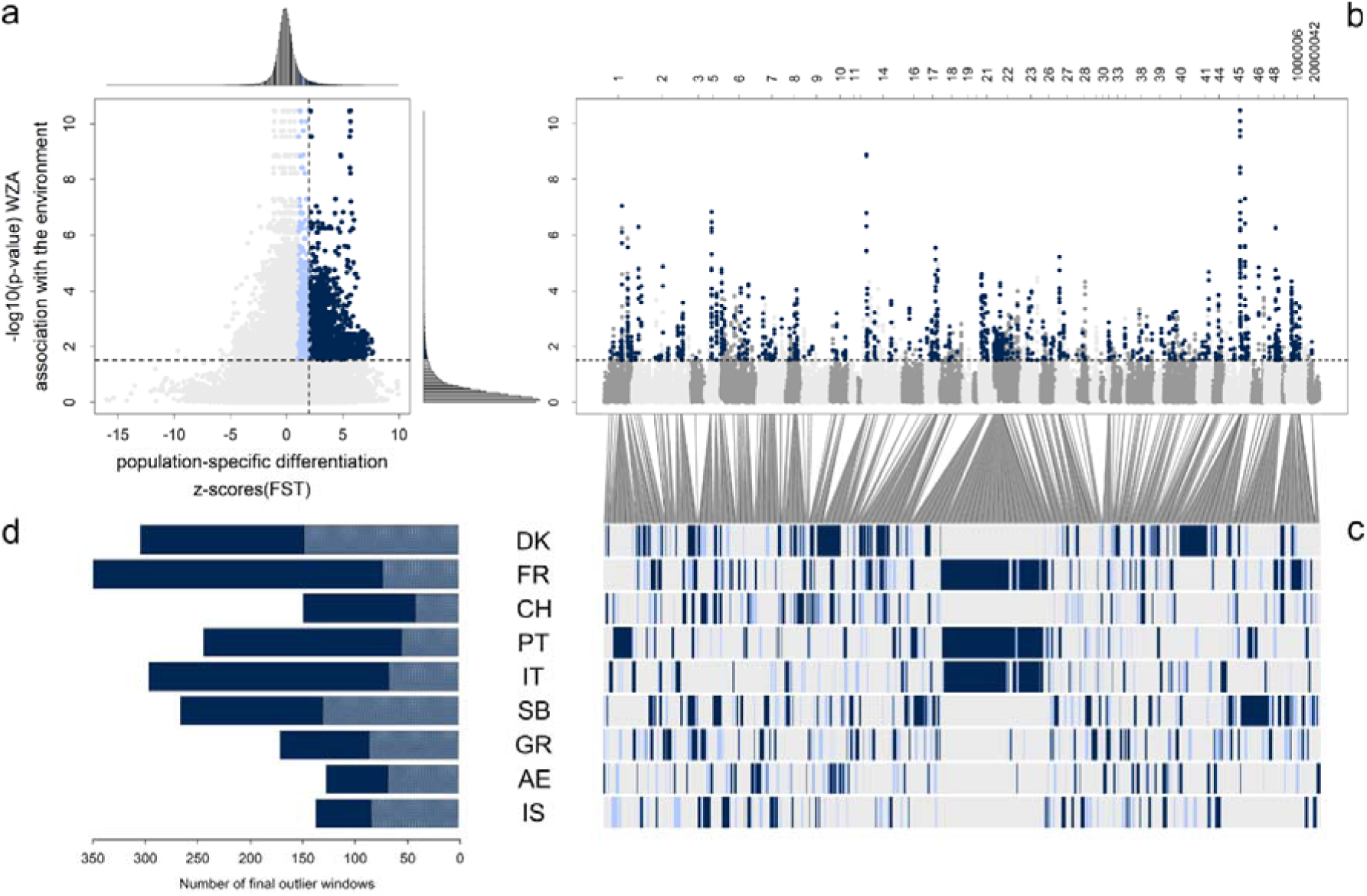
Genomic signatures of selection linked to climate in the European barn owls. (**a**) Scatterplot of genomic windows (100kbp each) across nine populations. Vertical and horizontal dashed lines are at 2 standard deviation from the mean of z-scores (population-specific FST) and z-scores (p-values) of WZA respectively. Each colour presents a class of windows. Dark blue dots represent outlier windows in both population-specific FST and WZA scans. Light blue dots are outlier according to WZA and have a population-specific FST higher than 1 standard deviation from the mean; (**b**) Genome-wide distribution of the WZA score. Dark blue dots corresponding to the windows identified in panel (a). A switch between light and dark grey represents a change in the scaffold. The names of all scaffolds are displayed on the upper x-axis. (**c**) Distribution of outlier windows in the different populations. Each row represents a population (DK: Denmark; FR: France; CH: Switzerland; PT: Portugal; IT: Italy; SB: Serbia; GR: Greece; AG: Aegean islands; IS: Israel). Each column is a window along the genome, coloured according to the classification in (a). (**d**) Barplot of outlier windows per population. Dark blue bars picture the outlier windows shared with at least one other population while white dashed bars picture the number of outlier windows unique to each population.

### Metabolic pathways associated with climate

To explore the functional implications of selection and climatic adaptation, we extracted a total of 550 genes from the 270 genomic regions that showed signatures of selection and variants associated with climatic variables. Among them, 324 genes were unique to a single population (Table S3). Within the 550 extracted genes, we detected significant enrichments in pathways related to cellular physiology (details of the Gene Ontology (GO) terms are provided in Table S4). We observed the same results when we used the population-specific list of genes from Greece, Israel and Serbia (Table S5). For the other populations, the GO enrichment analysis did not yield any significant results. Overall, the functional grouping of the 550 genes included functions such as immunity, locomotion, and anatomy in all the populations, with different genes in each population (Table S6).

### Climatic differences drive differential selection between Southern and Northern populations

Our analysis of the region with the highest signal of association with climate reveals that differential selection has led to marked genetic differentiation between populations at opposite ends of the temperature gradient. To explore the signal located on Super-Scaffold 45, we first examined climatic association values within this genomic region (Figure 3). Out of the 8 outlier windows showing the highest association with climate (red dots in Figure 3a-b), all were outliers in the population-specific FST scan for the Danish population, and two of them were also outliers in the Portuguese population, two populations at the opposite ends of the temperature gradient (first axis of Supplementary Figure 4). We assessed the extent of genetic similarity in this genomic region between the two populations by computing the population-pair FST, an estimate of the standardised mean kinship of individuals^31^. With this statistic, we detected a substantial reduction of the genetic similarity between the individuals from these two populations within the region harbouring the highest association with climatic conditions (red rectangle on Figure 3c). Consistently, we observed an increase of genetic dissimilarity by using a pairwise FST computed on an SNP basis between the two populations (Figure 3d). We examined the haplotypes within the highest section of the peak which showed a clear distinction between the Danish and Portuguese populations (Figure 3e).

**Figure 3.**
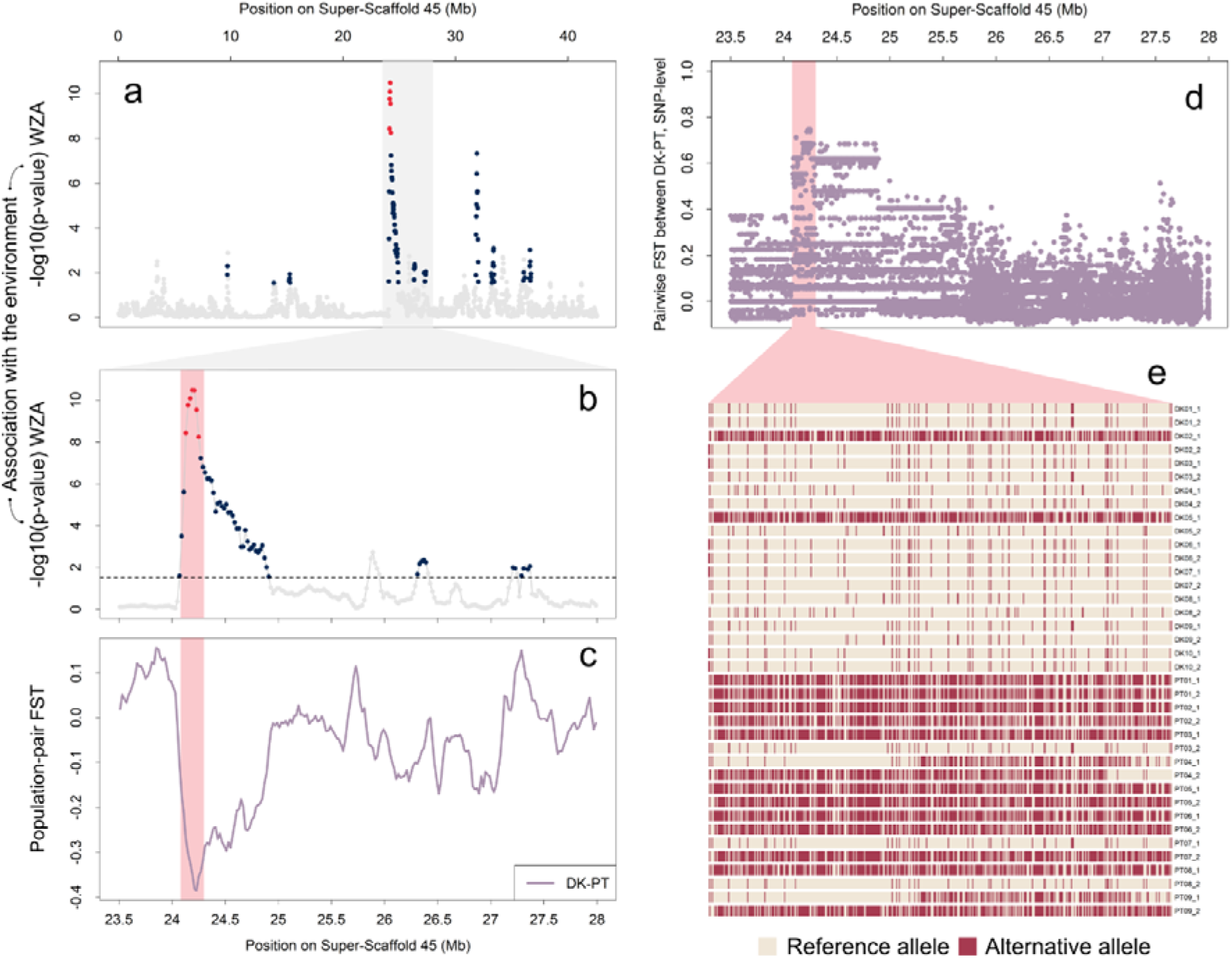
Divergent selection drives the strong climate-driven signal on Super-Scaffold 45. (**a**) WZA signal across Super-Scaffold 45 with the main peak of outliers highlighted in grey. Dark blue circles are outlier windows from WZA and population-specific FST. Red circles represent outliers with a -log10(p-values) higher than 8. (**b**) Zoom on this peak of the Super-Scaffold 45. (**c**) Population-pair FST in this region between two populations at the extreme of the first climatic axis in Figure 1b (namely Denmark and Portugal). Lower value indicates a higher divergence in this region compared to the rest of the genome. (**d**) Pairwise FST between Denmark and Portugal, computed on a SNP-basis. (**e**) Genotypes of each SNP within the highlighted region (red rectangle on panel (B), (C) and (D)), using phased haplotypes from Denmark and Portugal. Beige represents the reference allele, and red represents the alternative allele.

### A putative inversion linked to climatic adaptation in past refugia

We identified a long consecutive signal of climatic association shared between populations from France, Italy, and Portugal in the first half of Super-Scaffold 22 (Figure 2b-c; Figure 4a). We used the population-pair FST to assess whether the same genomic variants were shared in the three populations. We detected a higher genetic similarity between the three pairs of populations (FR-IT, FR-PT, and IT-PT) within a 14 Mb region at the beginning of the Scaffold than along the rest of the genome (Figure 4b). This coincides with a climatic convergence in former glacial refugia, where shifting conditions may have contributed to the shared adaptive signals (Supplementary Figure 9, 10, 11 and 12).

**Figure 4.**
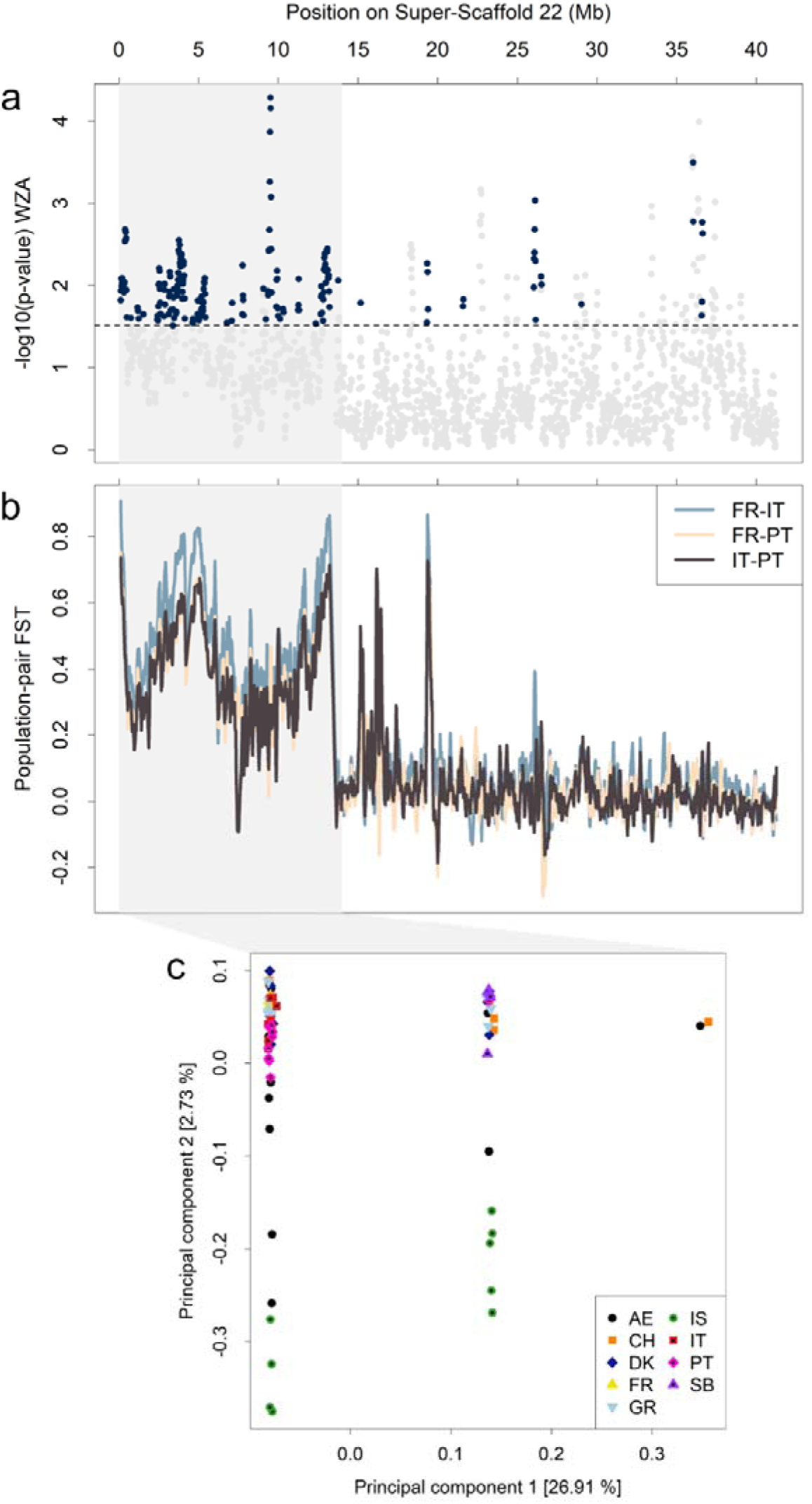
The shared signal in southern populations point to a putative inversion linked with local adaptation in the European barn owl. (**a**) WZA signal along Super-Scaffold 22 with the first 14 Mb highlighted in grey. Dark blue circles are outlier windows from WZA and population-specific FST. (**b**) Population-pair FST along the entire Super-Scaffold 22, higher value indicates a higher similarity in this region compared to the rest of the genome. As in (A), the first 14 Mb are highlighted in grey. Pairwise comparisons include France (FR), Italy (IT) and Portugal (PT). (**c**) PCA made on the first 14 Mb of the Super-Scaffold 22, (120,953 SNPs - DK: Denmark; FR: France; CH: Switzerland; PT: Portugal; IT: Italy; SB: Serbia; GR: Greece; AG: Aegean islands; IS: Israel).

To further investigate the genomic features of this region, we applied a PCA to the first 14 Mb of the scaffold, encompassing 120,953 SNPs, using data from all 74 individuals (Figure 4c illustrates the first two axes, explaining respectively 26.91 and 2.73% of the variation). Individuals were divided along the first axis into three distinct clusters, corresponding to the expected genomic pattern in case of a chromosomal inversion^32^. The leftmost cluster included more individuals than the one on the rightmost part of the x-axis. In comparison, the PCA conducted on the whole genome only separates the Portuguese samples from the rest along the x-axis and the Israel individuals from the rest along the y-axis (Figure 1d).

## Discussion

Climatic conditions and their past fluctuations are known to have shaped today’s distribution and genetic diversity of many species^1–3^. While genetic local adaptation is widely recognized as a key mechanism driving species persistence^6^, its role in shaping populations’ capacity to respond to future climatic shifts remains unclear. In this study, we integrated niche-based species distribution modelling (SDM^22^) with genome-wide analyses to investigate how climate-driven changes since the Last Glacial Maximum (LGM) have shaped the genetic composition and adaptive potential of the Western Palearctic barn owl. First, we showed that suitable conditions nowadays are more climatically diverse than the ones from the LGM^24–26^, exposing the individuals to heterogeneous and new selective pressures. Then, we identified genomic regions harbouring strong signals of selection and carrying variants associated with climatic variables. We observed signals of genomic adaptation in all the population we sampled across the species’ range, with a particularly strong signal shared among southern populations. Overall, our findings challenge the hypothesis that local adaptation primarily occurs at recolonising margins^33,34^, showing instead that genetically diverse populations such as past glacial refugia are a key source for adaptation^35^. We highlight that refugial populations, by maintaining standing genetic variation, serve as crucial reservoirs of adaptive potential, enhancing species’ ability to respond to future climatic changes^35–37^.

Species distribution modelling of the European barn owl based on its realized climatic niche suggests that the latter is broader nowadays than it was during the Last Glacial Maximum (LGM). We used the Maximum Entropy method^38^ for fitting the SDM and project it onto past climatic conditions to compare the extent of suitable habitat under current versus historical climates. Our results indicate that some contemporary climatic conditions along the Atlantic coast, up to the UK, were already present 20,000 years ago. This supports the hypothesis that, as temperatures rose at the end of the LGM, favourable conditions shifted toward northwestern Europe, facilitating barn owl range expansion^39^. However, we also identified newly emerged suitable climatic conditions that were absent during the LGM in recolonised Central and Eastern Europe as well as in the glacial refugia where conditions changed for resident populations. All these results suggest that the realised climatic niche of the barn owl was contracted during the Last Glacial Maximum and extended with the subsequent warming. We advance that genetic adaptation played a significant role in this process and further explored this hypothesis.

By combining population-specific FST scans^27,28^ to identify genomic regions under selection with genome-environment association approaches, we found genomic evidence of local adaptation to humidity and temperature in all populations. Genes among selected regions showed significant GO term enrichment in several populations, but we found no clear link to climate adaptation. However, the functional clustering revealed a concentration of genes related to immunity, with GO terms such as “immune system process” and “immune response” (Table S6). This finding aligns with previous research showing that pathogens and infectious diseases exert strong selective pressures on both humans and birds^40,41^. Moreover, temperature and rainfall significantly influence pathogen community composition^42,43^. Supporting this connection between climate and immune genes, O’Connor et al. (2020) demonstrated that MHC-I genes in birds vary in diversity according to humidity^44^. GO terms related to anatomy and growth were also found across population, encompassing several dozens of genes linked to “anatomical structure morphogenesis”, “regulation of the developmental process” and “growth” (Table S6). This result was consistent with Bergmann’s rule which predicts that body sizes of warm-blooded vertebrates negatively correlate with temperature, leading to smaller body sizes in warmer climates^45,46^. Overall, our results suggest an implication of these genes in local adaptation to various climates, but further work is needed to identify and confirm their precise role.

A closer look at the regions of the genome with many variants linked to the environment and with strong traces of selection revealed a wide range of mechanisms shaping the barn owl’s local adaptation, from opposite directional selection to recurrent evolution.

The region located on the Super-Scaffold 45 showed a clear signal of association with climatic variables. In this region, population-specific FST highlights the high differentiation between the Portuguese and Danish populations, which lie at the opposite ends of the temperature gradient (Supplementary Figure 4). This, along with their distinct haplotypic structure (Figure 3), suggests that different alleles or haplotypes are under selection in these populations^47^. A closer look at the genes present in the region did not reveal any genes that can easily be linked to temperature adaptation and further work should confirm whether this region plays a role in thermal adaptation.

Our results also revealed a strong and consistent genomic signal of selection linked to climate across all individuals from three populations, one in a region with a climate similar to that of the LGM (France) and two in past glacial refugia (the Iberic and Italian peninsulas). The size of the region involved; the pronounced pattern of genetic differentiation compared to the rest of the populations as well as the cluster patterns found by our local PCA strongly suggest the presence of an inversion in this region^32^. Supported by the growing body of literature linking inversion and adaptation^48–50^, we suggest that this inversion is adaptive to climatic conditions. Briefly, considering the genomic disruption that breakpoints can cause, as well as the fact that nearly suppressing recombination will prevent purging of deleterious alleles in the inverted haplotype, it is unlikely for a newly appeared inversion to persist and spread in a population^51^. However, by bringing adaptive alleles together and averting maladapted gene flow by blocking recombination, an inversion can promote local adaptation and thus be favoured by selection^50,52,53^.

The history of this inversion remains to be explored. The pattern of selection is shared between the two peninsulas that hosted the species during the LGM. Considering the isolation of the two populations at the time and the reduced connectivity nowadays (see Figure 4 in Cumer et al., 2021 for details)^27^ we suppose that this inversion predates the LGM, and that independent and recurrent selection drove the increase of frequency of this haplotype independently in these two populations. Further work should formally test this hypothesis.

A key question is the origin of adaptive alleles and given the pattern of genomic traces of local adaptation we observed, we propose that standing variation played a crucial role in this process^54,55^. One possibility for local adaptation to occur is de novo mutations bringing new favourable alleles in the different populations^54,56^. Considering the low mutation rate^57,58^ and the even lower probability of a mutation being both advantageous and occurring in the right environmental context^59^, this mechanism is unlikely to fully explain the extent of genomic regions involved in local adaptation (Figure 2). A more parsimonious explanation relies on adaptation to pre-existing genetic variation^55^. In this scenario, allele sorting from standing genetic variation would have driven adaptation^60^. This occurs when previously neutral or mildly deleterious alleles become advantageous following shifts in selective pressures, such as habitat colonisation or climate change^60^. This scenario implied that within glacial refugia, a pool of segregating alleles was maintained in sufficiently large populations^61^. These alleles were then sorted through space by local adaptation, with different environments (biotic and abiotic) acting as a filter on deleterious and/or less advantageous alleles. This whole process has been reported in species as diverse as insects (Drosophila melanogaster^62^), birds (Darwin’s finches^63^) mammals (humans^59,64^) and fish (sticklebacks^65^). We did not explicitly test whether local adaptation in barn owls resulted from de novo mutations, allele sorting from standing variation, or a combination of both. Therefore, additional work, focusing on the date of mutations^66^ as well as the timing of selection^67^, should confirm our results. However, the pattern of local adaptation that we identified in populations from past glacial refugia already provides insight into how standing variation fuels adaptation during climatic fluctuations.

The climatic variation since the Last Glacial Maximum provides a good framework to gain knowledge on where and how populations adapt to climate change. A wide body of literature explored many aspects of what happens in populations that expanded into newly available habitat, from dispersal limitation^68,69^ to the genetic load accumulated during an expansion^70–74^, and how these mechanisms interact with local adaptation^75–78^. However, the fate of refugial populations remains, to our knowledge, understudied. In this work, we first demonstrate that the climatic conditions experienced by these populations have changed over time, creating ongoing pressures for adaptation. Our findings suggest that refugial populations are not only well adapted to their current environments but are also continuing to adapt. Based on our results, we propose that adaptation in refugial populations is a recurrent and dynamic process. As the climate continues to shift, the species’ optimal niche moves northward, and southern populations will be among the first to experience novel climatic conditions. These populations often harbour the highest levels of genetic diversity^17^, which likely confers them the greatest adaptive potential in the face of climate change. This underscores the importance of conserving these genetically diverse populations, whose adaptive capacity may be critical for the long-term resilience of the species. While we acknowledge that the pace of current climate change is far more rapid compared to shifts since the Last Glacial Maximum, our study las the groundwork for a deeper of the genomic foundations of adaptative potential. Future research will be essential to unravel the mechanisms that enable species to persist and thrive in rapidly changing environments.

## Material and methods

### Ecological modelling

#### Species distribution modelling

We first conducted species distribution modelling (SDM^22^) to identify suitable areas for the species during the Last Glacial Maximum and nowadays. We fitted SDM using the Maximum Entropy modelling software (MaxEnt^38,79^, v.3.4.3), a presence-only based procedure in a similar approach as the one described in Cumer et al. (2022)^15^.

First, we extracted 19 bioclimatic variables at a 5 arc-minute (∼9.3 km at the equator) resolution from the WorldClim 1.4 database^80^ using the rbioclim R package^81^. To avoid redundancy between variables, we removed variables with a correlation equal to or higher than 0.8^82^, leading to a set of 7 uncorrelated climatic variables: Mean Diurnal Range (Bio2), Min Temperature of Coldest Month (Bio6), Temperature Annual Range (Bio7), Mean Temperature of Wettest Quarter (Bio8), Precipitation Seasonality (Bio15), Precipitation of Driest Quarter (Bio17), and Precipitation of Coldest Quarter (Bio19).

We performed the analysis using the dismo R package^83^ v.1.3-14 on R v.4.3.2. To determine which combination of parameters optimised the model without over-complexifying it^84^, we built models with linear, quadratic, and hinge features, using a range of regularisation multipliers from 1 to 5 (as recommended in Warren & Seifert, 2011^85^). A quadratic feature with a regularisation multiplier of 1 yielded the lowest AIC and was chosen for further modelling. We ran a total of 100 quadratic MaxEnt models (with a regularisation multiplier = 1), omitting 25% of the data during training to test the model and using 10’000 background points randomly sampled across the study area. For each model, we randomly sampled 1,000 presence points from the IUCN distribution map^86^. We evaluated the predictive performances of the models by assessing the area under the curve (AUC) of the receiver operating characteristic (ROC) plot of the test data^87^. All models had an AUC between 0.756 and 0.807 (see Supplementary Figure 13), thus classified as fair to good according to Li et al. (2020)^88^.

We projected the 100 models to the present climatic conditions and to the climatic conditions from the Last Glacial Maximum (about 20’000 years ago), also extracted from the WorldClim 1.4 database (scenario CCSM from PMIP2^80^) at 5 arc-minute resolution. We used the “maximum training sensitivity plus specificity” (MaxSSS) threshold, as recommended for presence-only data^89^, to transform the projected output from the models into binary suitability maps (0 unsuitable, 1 suitable). Finally, we averaged the values among replicates and retained cells as suitable only if they were so in at least 90% of the models. To avoid model extrapolation when projecting the models in the past, we used the Multivariate Environmental Similarity Surface (MESS) approach^90^ to identify and discard areas from the past with climatic conditions absent from those in the calibration data.

#### Identification of newly suitable conditions absent from the LGM

Then, we determined whether currently suitable areas display climatic conditions different from those found during the LGM. To do so, we extracted values of the 7 climatic variables used in the SDM (see Species distribution modelling section for details) at every continental cell of the studied area for both past and present and performed a Principal Component Analysis (PCA) to represent the environmental space. We retained the first three principal components that explained up to 91.50% of the climatic variance. In this environmental space, we generated a “multidimensional climatic space”, representing the entire range of climatic conditions suitable for the barn owl, by combining past and present suitable pixels within this PCA space. We then characterised the past suitable conditions for the species by creating a polygon around the LGM suitable conditions, using the alphashape3d R package (α = 0.5, keeping all the data^91^). Next, we classified the suitable conditions at current time as located inside or outside the polygon, thus respectively present or absent from the range of LGM suitable conditions.

#### Estimation of the shift of climate within suitable areas

To quantify shifts in climatic conditions between LGM and now, we identified cells that were suitable both during the LGM and present time (Supplementary Figure 9) and computed the Euclidean distance between their past and present positions in the multidimensional climatic space. We then rescaled the distance values from 0 to 1 (Supplementary Figure 10).

Because we discovered a strong signal of selection shared between the two glacial refugia populations (see Results section), we investigated how the climate has changed since the LGM in the two peninsulas to assess whether the climate converged or diverged in these two locations. We extracted values of the 7 climatic variables used in the ecological modelling for the LGM and today (see Species distribution modelling section) at the GPS coordinates for the Italian and Portuguese samples. We projected these conditions inside the multidimensional climatic space (Supplementary Figure 11). Then, we computed the pairwise Euclidean distances between all sampling localities of the two peninsulas during the LGM and the present time (8 (IT) x 8 (PT) pairwise distances at both time points). This way, we quantified the climatic difference between the Italian and Portuguese sampling localities at the two different time points (Supplementary Figure 12).

### Individual barn owl sampling

#### Biological samples

This study took advantage of the datasets of European barn owls previously published by Cumer et al. (2022), Machado, Cumer, et al. (2021), and Machado, Topaloudis, et al. (2021)^15,27,92^. We retrieved the whole genome sequences from the Sequence Read Archive (SRA - Bioprojects PRJNA700797, PRJNA727915, and PRJNA727977, Table S1).

#### Genetic data preparation

We performed the read mapping, variant discovery, and variant filtering following Cumer et al. (2022)^15^. In brief, we mapped raw reads to the reference barn owl genome (GenBank accession GCA_018691265.1^39^) with BWA-MEM v.0.7.15^93^. We performed Base quality score recalibration (BQSR) in GATK v.4.1.3 using high-confidence calls described in Cumer et al. (2022)^15^. We called variants with GATK’s HaplotypeCaller and GenotypeGVCF v.4.1.3 from the recalibrated BAM files. We filtered genotype calls using a hard-filtering approach as suggested for nonmodel organisms, using GATK and VCFtools^94^. Details of technical filtration can be retrieved from Cumer et al. (2022)^15^.

To prevent some alleles from being over-represented in the dataset (relatedness statistic based on the method of Manichaikul et al., 2010^95^, implemented in VCFtools v0.1.14^94^), we identified pairs of individuals with a relatedness higher than 0.1. We removed one of the two individuals for each identified pair, leading to a dataset of 74 individuals (Table S1).

We then excluded genomic regions with uncertain SNP calling by removing regions of the genome where our ability to confidently map reads is limited (i.e., a “mappability” mask). To achieve this, we followed the procedure documented at lh3lh3.users.sourceforge.net/snpable.shtml. In summary, we divided the reference genome into reads of 150 base pairs (bp) with a sliding window of 1 bp. These artificial reads were then mapped back to the reference using BWA-MEM v.0.7.17. Regions of the sequence where less than 90% of the reads mapped perfectly and uniquely were discarded by excluding variants using a bed file in VCFtools v0.1.14. We retained only bi-allelic SNPs, removed loci with more than 5% missing data, and excluded from the analysis 15 scaffolds (out of 60) showing less than 1000 SNPs and one extra sexual scaffold.

At the end of the filtration process, we retained a set of 12,309,943 SNPs from 39 scaffolds, genotyped in 74 European barn owls (10 individuals from the Aegean Islands (AE), 10 from Denmark (DK), 4 from France (FR), 9 from Greece (GR), 9 from Italy (IT), 10 from Israel (IS), 9 from Portugal (PT), 5 from Serbia (SB), and 9 from Switzerland (CH)).

To explore the genome-wide variations of the Western palearctic populations and check its concordance with previous results, we conducted a PCA with the SNPRelate R package^96^ using this entire set of SNPs and individuals. Additionally, we assessed genetic diversity among the 9 sampled populations following the procedure of Cumer et al., (2022)^15^. Briefly, we identified the number of polymorphic sites, private alleles, rare alleles and the whole genome population-specific FST for each population independently. To account for differences in sample sizes (ranging from 4 to 10), we randomly sampled 5 individuals from each population - except for FR and SB - and calculated these diversity estimates on the resulting subsets. This resampling process was repeated 10 times, and we reported the mean and standard deviation of the diversity estimates.

#### Phasing process and evaluation

We performed the phasing and imputation of the individuals’ genotypes in two steps. First, we conducted a read-based phasing of each individual using WhatsHap v1.0^97^. During this step, we reconstructed haplotypes based on the mapped sequencing reads covering multiple variants. Between the two filtering steps, we applied a Minor Allele Frequency (MAF) filter to ensure it was higher than 5% using VCFtools v.0.1.14^94^, thereby removing rare alleles from the dataset that could influence the second round of phasing, resulting in a dataset of 4’689’284 SNPs. Then, we conducted the complementary round of phasing with ShapeIt4 v4.1.3^98^ with default parameters. The latter uses a statistical approach to infer individuals’ haplotypes based on the population genotypes^99^ and incorporates the phase information from the read based phasing.

To evaluate phasing performance, we calculated the switch error rate (SER) of the phasing generated by ShapeIt4 for each individual^100^. For each individual, we conducted a statistical phasing using ShapeIt without considering the read-based phasing from WhatsHap for the focal individual. Subsequently, we compared this phasing to the “true” local phasing, inferred from the read-based approach (WhatsHap). We estimated the switch error rate between both sets of phasing using the switchError code (available at https://github.com/SPG-group/switchError). Among the 74 phased individuals, the mean error rate was 2.23 * 10^-4^ and none exceeded 0.7% (Supplementary Figure 14).

### Detection of traces of selection in each population

We computed a single summary statistic to identify genomic regions potentially under selection: population-specific FST, using the hierfstat R package^101^. The statistic was calculated across the genome in overlapping windows of 100 kbp with 20 kbp steps for each independent population. Only windows containing at least 250 SNPs, corresponding to two standard deviations below the mean, were included in the analysis.

To identify outlier windows (regions with extreme values of positive population-specific FST), we first transformed the statistic into Z-scores by subtracting the population mean from each estimate and dividing it by the standard deviation for each population independently. We then combined Z-scores from all populations and considered a window as an outlier in each population if its Z-score for population-specific FST was equal to or higher than 2 standard deviations from the mean of the merged Z-scores. This approach allowed us to focus on regions exhibiting an excess similarity in the population compared to the rest of the genome (high population-specific FST). For further details about this method, refer to Cumer et al. (2022)^27^. We computed the statistic on the set of 12,309,943 SNPs, unfiltered for Minor Allele Frequency (MAF). To ensure that rare alleles did not influence population-specific FST, we also calculated this statistic on the filtered variants from MAF (set of 12’309’943 SNPs). The high consistency between the two estimates (with and without MAF filtering), as depicted in Supplementary Figure 15, supported our decision to retain the statistic computed on the unfiltered variants set (12,309,943 SNPs).

### Genotype-Environment Association

#### Redundancy Analysis

We independently conducted a Genotype-Environment Association (GEA) analysis to assess the relationship between the genotypes of the barn owl and their surrounding environment for all populations simultaneously.

Based on the GPS coordinates of the 74 samples (Table S1), we extracted values for the same 7 climatic variables as those used in the species distribution modelling (Mean Diurnal Range (Bio2), Min Temperature of Coldest Month (Bio6), Temperature Annual Range (Bio7), Mean Temperature of Wettest Quarter (Bio8), Precipitation Seasonality (Bio15), Precipitation of Driest Quarter (Bio17), and Precipitation of Coldest Quarter (Bio19); see Species distribution modelling section for details). We conducted a Principal Component Analysis (PCA) on these climatic data to assess the level of climatic dissimilarity experienced by the barn owls sampled in this study nowadays. We retained the first three principal components, explaining 94.97% of the climatic variance.

We then associated variants with genomic information using Redundancy Analysis (RDA) with the vegan R package^29,102^. This method relies on a multiple linear regression of the observed genotypes on a set of abiotic or biotic predictors. The expected genotypes based on the model (also called fitted values) are then extracted and used as input for a PCA called RDA space. The projection of the principal axes and components in this RDA space allows the detection of the SNPs that contribute the most to the RDA axis and whose allelic frequency might be putatively driven by the explanatory variables^29^. We used the imputed genotype matrix (phased set of 4’689’284 SNPs) as the response matrix and the bioclimatic variables extracted at each sampling locality for the multiple linear regression.

To evaluate the significance of the relationship between genotypes and climatic variables, we performed a permutation test^103^. In brief, we computed a test statistic (F-statistic) from the regression using the true data. Afterwards, we carried out 999 additional regressions on permuted rows of the response data (i.e., the genotype matrix), allowing us to establish the empirical null distribution of the statistics, to which we compared the observed statistic^103^. From this test, we found a significant relationship between the environmental variables and genetic components (Table S7). To select the number of RDA axes to retain, we also performed a permutation test for each axis (n = 100) by following the procedure given by Borcard et al. (2011)^103^. Applying this method, we kept the first five RDA axes, explaining 80.41% of the constrained variance (Table S8). To detect loci that strongly contribute to the individual’s discrimination in the RDA space (outlier loci), we followed the procedure described in Capblancq et al. (2018)^104^: we computed Mahalanobis distances between each locus and the centre of the RDA space using the previously retained five axes. P-values were adjusted for the false discovery rate (FDR) by computing q-values using the qvalue R package^105^. We considered a SNP as an outlier if its q-value was less than 0.1, following Capblancq et al. (2018)^104^.

#### Weighted-Z analysis

To improve the robustness of our GEA approach, we decided to transform the SNPs p-values from the Redundancy Analysis into window-based statistics through the Weighted-Z analysis (WZA) proposed by Booker et al. (2021)^30^. This method takes as input individual p-values from any SNP-based GEA approach and calculates a weighted-Z statistic for a given genomic region. To do so, it transforms the p-values of the focal window into z-scores and computes the weighted-Z statistic using the equation provided by Booker et al. (2021)^30^, which considers the variation in the number of SNPs among windows along the genome.

We computed the weighted-Z statistics on the same windows as population-specific FST. Since WZA does not support overlapping windows, we split the window set based into five sets of non-overlapping windows. We ran 5 separate weighted-Z analyses, one with each input file, and merged the outputs to obtain the final one. We considered a window as an outlier when the -log10 of its p-value was equal to or higher than 2 standard deviations from the mean of all windows (equivalent to a p-value of 0.03 assuming a normal distribution).

### Genomic signal of local adaptation to climate

#### Concordance between the genome scans and landscape genomics

Because the study aimed to detect traces of selection linked with local climatic conditions, we considered the final list of outlier windows as the overlap between the outlier set from the genome scans (see Detection of traces of selection in each population section) and the one from the Genotype-Environment Association analysis (see Weighted-Z Analysis section). Based on the annotated barn owl genome (GenBank Assembly Accession: GCA_018691265.1), we considered genes that partially or fully overlapped the final list of outlier windows as potentially involved in local adaptation to climate and extracted a list of genes for each population.

#### Gene Ontology Enrichment

We conducted Gene Ontology Enrichment (GOE) analyses using ShinyGo v.0.8^106^ to assess which biological pathways the genes located in the final list of outlier windows could be involved in and whether we could link some to local adaptation to abiotic conditions. We performed GOEs for each set of genes from each population separately and one additional GOE using all genes detected in at least one population. As a baseline set of genes against which we compared the observed enrichment signature, also called the background list, we used all the genes annotated in the barn owl genome that could have been detected with our 52’429 non-overlapping windows from the genome scans or WZA. For each analysis, we used the default pathway databases, namely KEGG (Kyoto Encyclopedia of Genes and Genomes), as well as the GO Biological Process database.

### Exploration of signals specific to some populations

#### Population-pair FST on Super-Scaffold 45

We identified a strong environmental association in Denmark and Portugal, two populations at the opposite of our environmental gradient, with strong population-specific FST in this region in each of them. To know whether populations have the same genetic variants or opposite ones, we assessed the level of genetic (dis)similarity between pairs of populations on this part of the genome. To do so, we computed a population-pair FST, described in detail in Goudet & Weir (2023)^31^, using the hierfstat R package^101^. In brief, for every pair of populations, we calculated the average kinship among pairs of individuals and standardised it by the average kinship between all populations using the same dataset and windows employed for population-specific FST analysis. A schematic example is provided in Supplementary Figure 16 for the Denmark - Portugal population pair.

#### Pairwise FST at the SNP level

To calculate a pairwise FST at the SNP level^31^ and confirm the signal obtained through population-pair FST, we used the fs.dosage function of the hierfstat R package^101^. We computed this pairwise FST on the entire Super-Scaffold 45 between Denmark and Portugal using the same dataset as for population-specific FST or population-pair FST.

#### Local PCA on Super-Scaffold 22

As we detected a strong signal of selection related to climate in the first half of Super-Scaffold 22, we decided to investigate the genetic architecture of this region. To do so, we conducted a PCA on the first 14 Mb of the scaffold (120,953 SNPs) using all individuals, using the SNPRelate R package^96^.

## Acknowledgements

We thank Olivier Delaneau for his advice on the phasing of the individuals and its evaluation. We thank Marc Robinson Rechavi for his advice on the Gene Ontology Enrichment analysis. This work was supported by the Swiss National Science Foundation with grants 31003A_179358 to JG.

## Data and code accessibility

The data underlying this article are available in the GenBank Sequence Read Archive Database at https://www.ncbi.nlm.nih.gov/sra, and can be accessed within Bioproject PRJNA700797, Bioproject PRJNA727915 and Bioproject PRJNA727977. Code used in this article is available at https://github.com/hugocrvl/TytoAlbaLocalAdaptation. Additional data can be provided by the corresponding authors.

## Authors contribution

HC, TC, and JG designed this study; TC called the variants; AT produced the mappability mask; HC conducted the analyses with the help of TC and suggestions of all authors; FC and AG advised on the SDM; HC and TC led the writing of the manuscript with inputs from all authors.

