## Supplementary Material for "Genetically diverse populations hold the keys to climatic adaptation: a lesson from a cosmopolitan raptor"

### Supplementary Figures

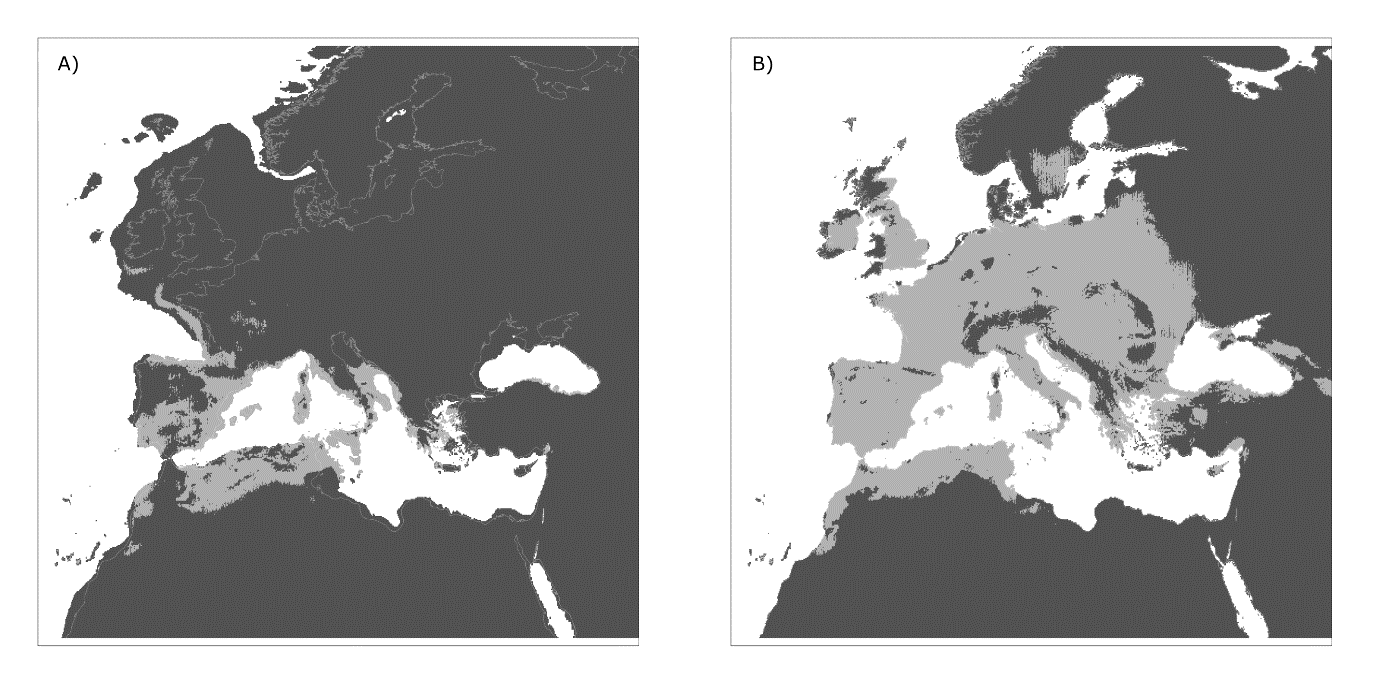

Figure S1 – Species distribution model of the Afro-European barn owl (Tyto alba) based on climatic variables, projected into the past (Last Glacial Maximum (20 kya)) - panel (A) - and present condition - panel (B). Locations in light grey were classified as highly suitable in 90% of the models. Below that threshold cells were considered unsuitable (darkest grey shade on the graph). The present coastline is outlined in white in all graphs.

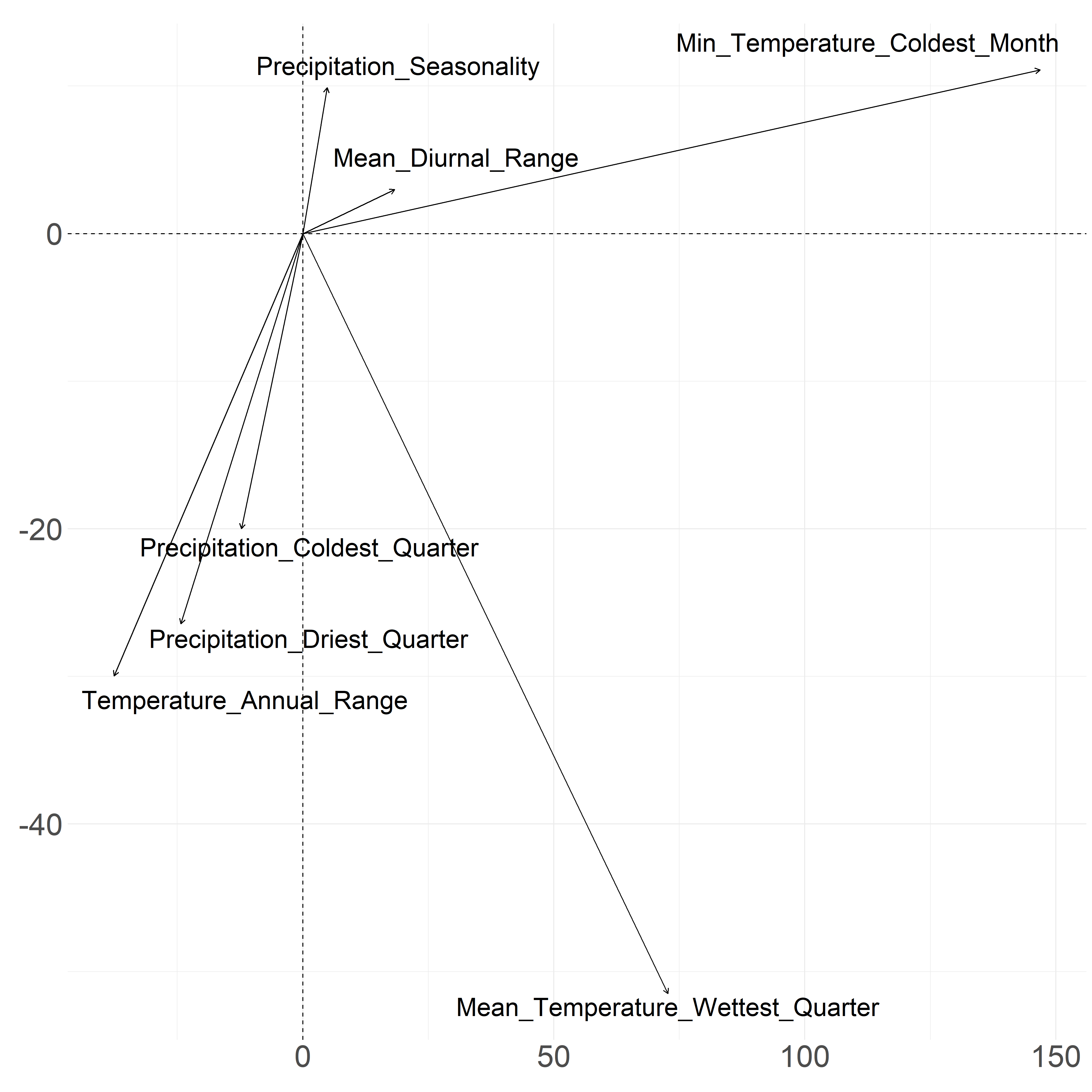

Figure S2 - PCA on bioclimatic variables extracted on the entire study area in the past (LGM) and present time. The first principal component is represented on the x-axis; The third principal component on the y-axis.

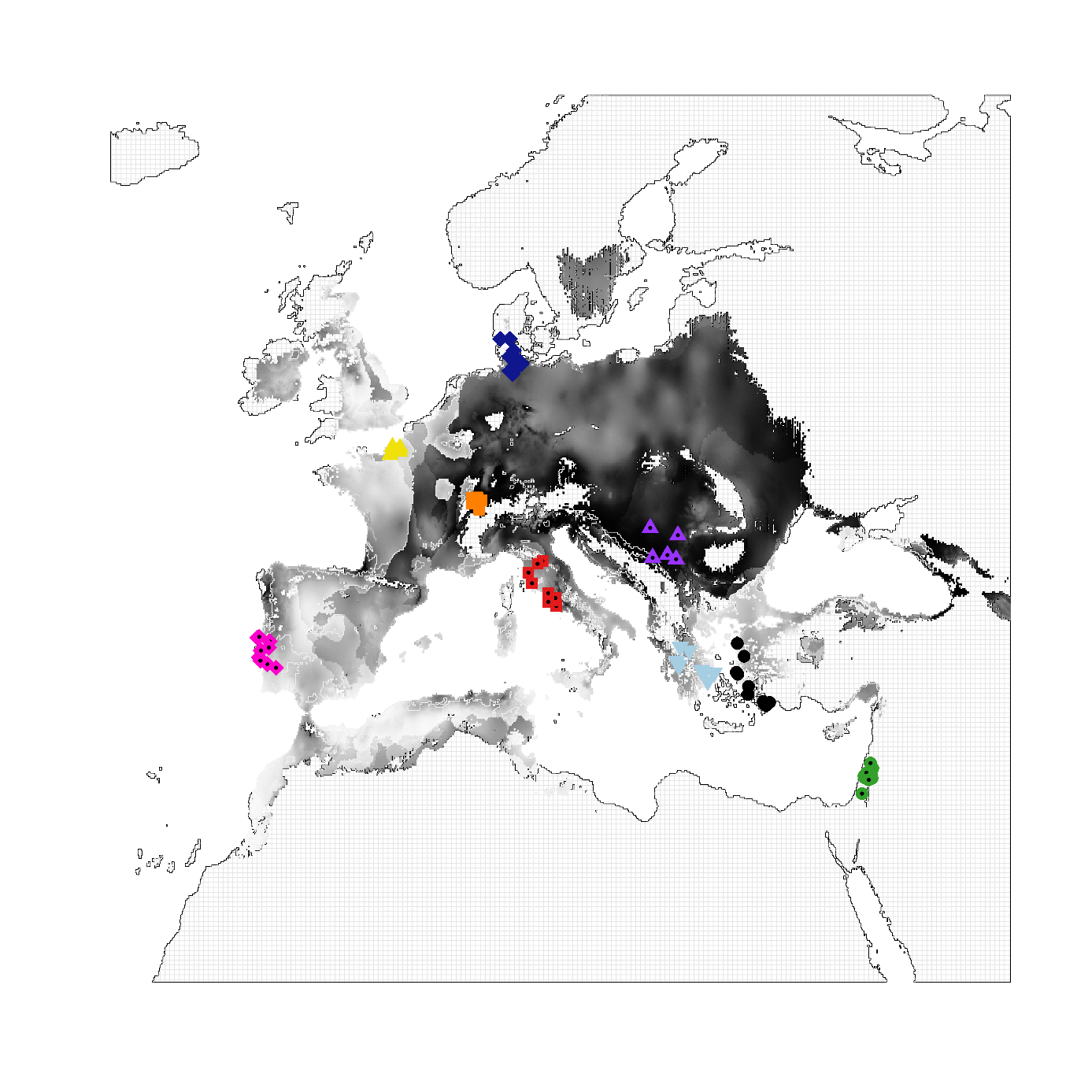
Figure S3 – Projection at the European scale of the contribution of the third principal component (PC3) from the PCA made on bioclimatic variables. The dark to white gradient signifies the contribution intensity; the brighter the colour, the higher the contribution of PC3 at this geographical location. Symbols represent sampling coordinates of individuals from 9 different populations, with legends being the same as in Figure 1.

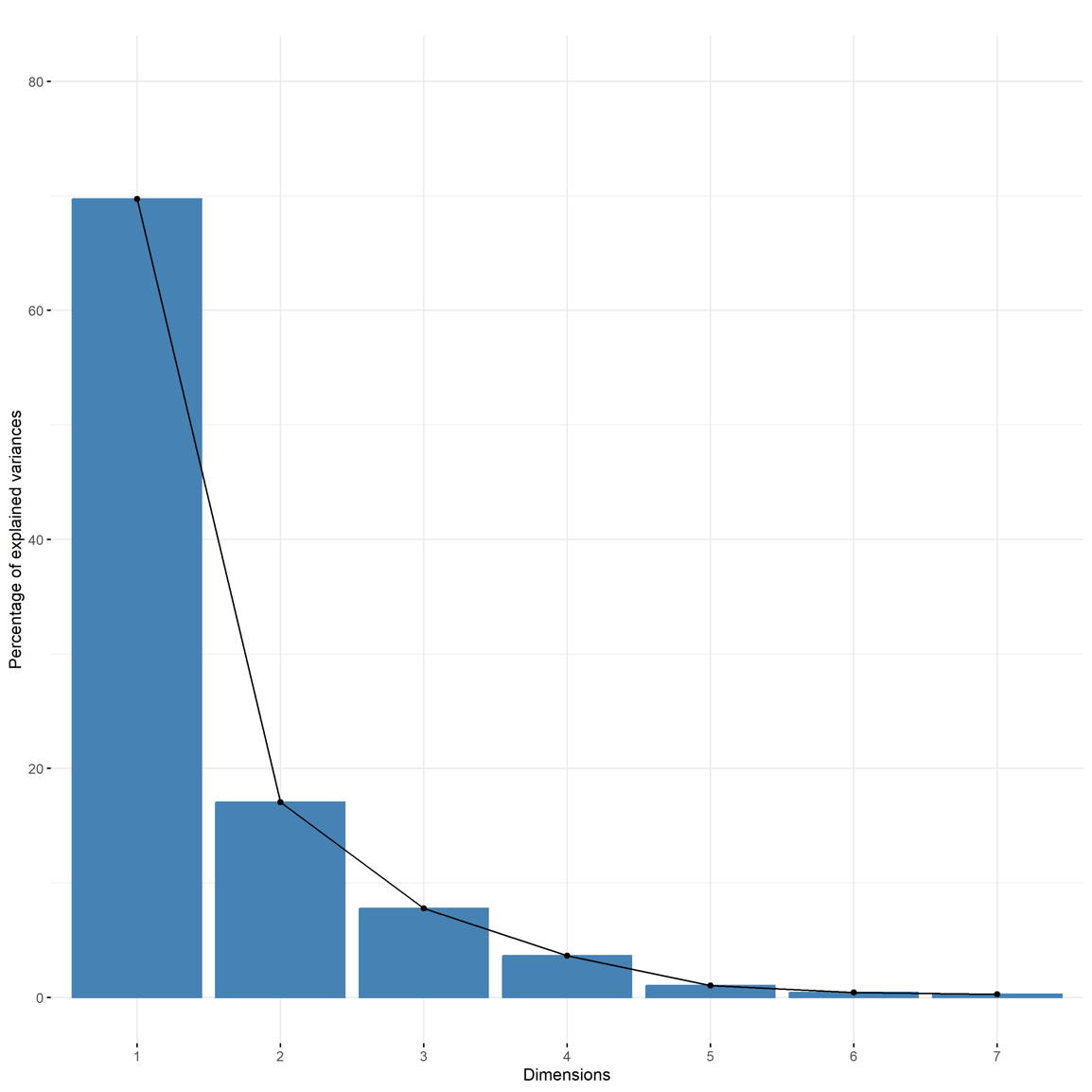

Figure S4 - Percentage of variance explained by the principal components from the PCA made on the bioclimatic variables extracted at the 74 sampling coordinates. We retained the first three principal components as they explained 94.97 % of the variance.

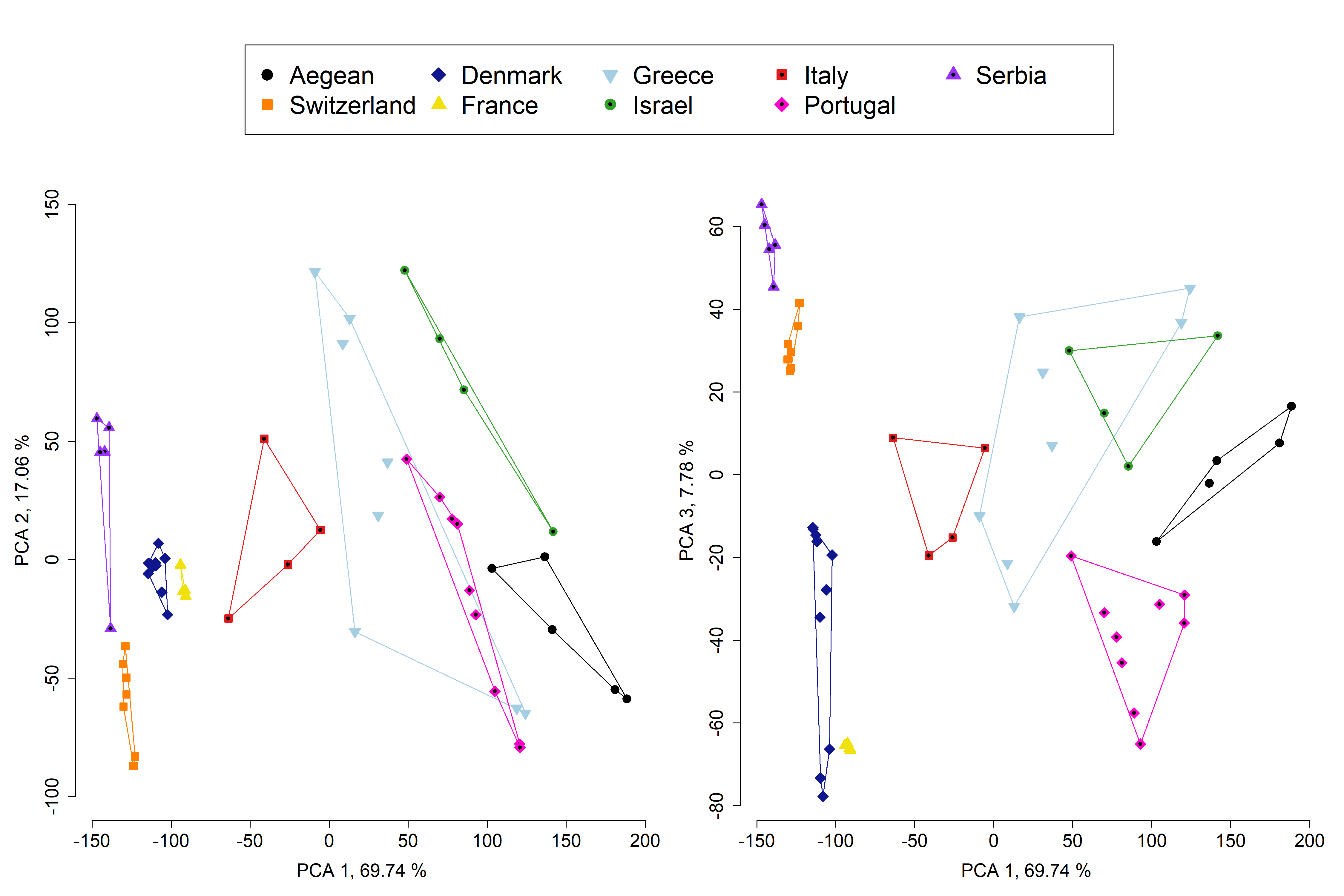
Figure S5 – Heterogeneity of the climatic conditions encountered by the 74 sampled barn owls. PCA was applied to bioclim values extracted from the 74 GPS sampling localities of the European barn owl used in this study. Climatic variables were the same as the ones used in the SDM (See Species distribution modelling section). The first three principal components explained 94.5 % of the climatic variance. The first axis was mostly driven by driven by temperature variables (specifically Mean Temperature of the Coldest Month (bio6), Temperature Annual Range (bio7), and Mean Temperature of the Wettest Quarter (bio8)). The second axis was mostly driven by precipitation variables (particularly Precipitation Seasonality (bio15), Precipitation of the Driest Quarter (bio17), and Precipitation of the Coldest Quarter (bio19)). The third axis was driven by bio7 (temperature annual range) and bio8 (mean temperature of the wettest quarter).

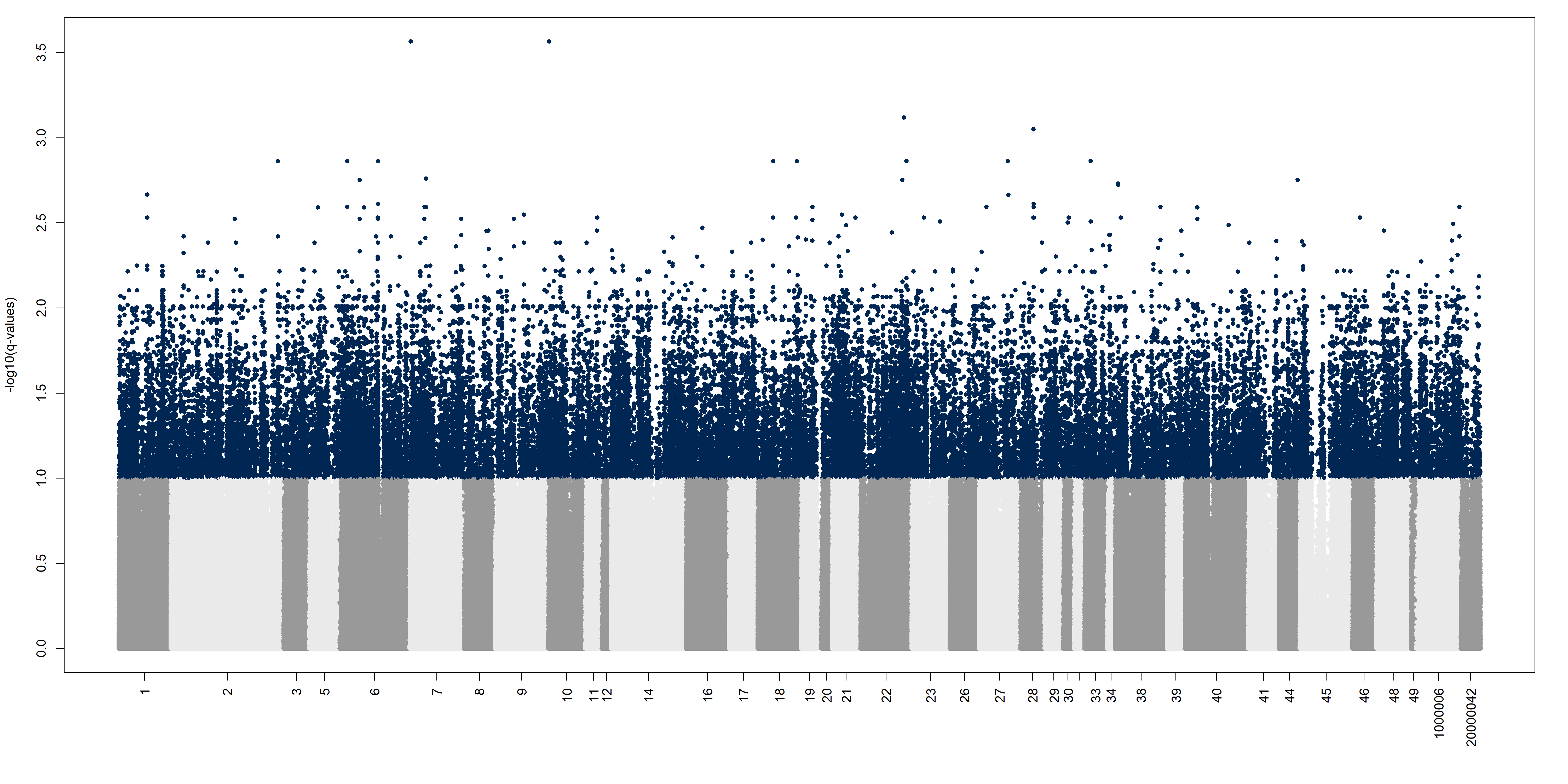

Figure S6 – Manhattan plot of the results from the Redundancy Analysis. We computed Mahalanobis distances between each locus and the centre of the five RDA axes. P-values were adjusted for the false discovery rate (FDR) by computing q-values. A value is considered an outlier (coloured here in dark blue) if its q-value was less than 0.1. A switch of colour between dark and light grey represents a change in the scaffold, named on the x-axis.

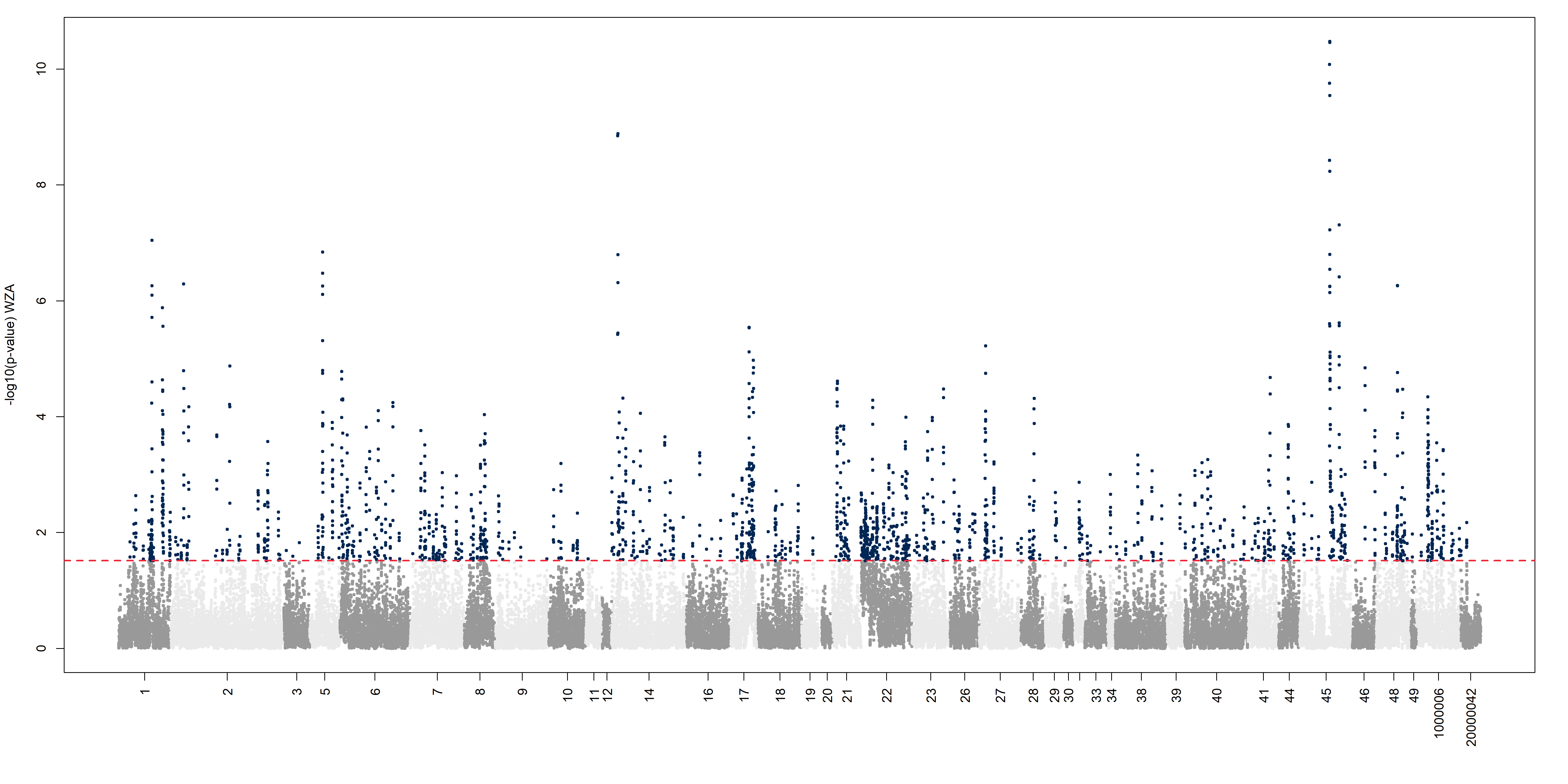

Figure S7 – Manhattan plot of the signal from the Weighted-Z analysis. Signal from the RDA were transformed on a window-basis statistic. We used the same windows as the ones from population-specific FST. A window was considered an outlier (coloured in dark blue here) if the -log10 of its p-value was equal to or higher than 2 standard deviation from the mean (equivalent to a p-value of 0.03). A switch of colour between dark and light grey represents a change in the scaffold, named on the x-axis.

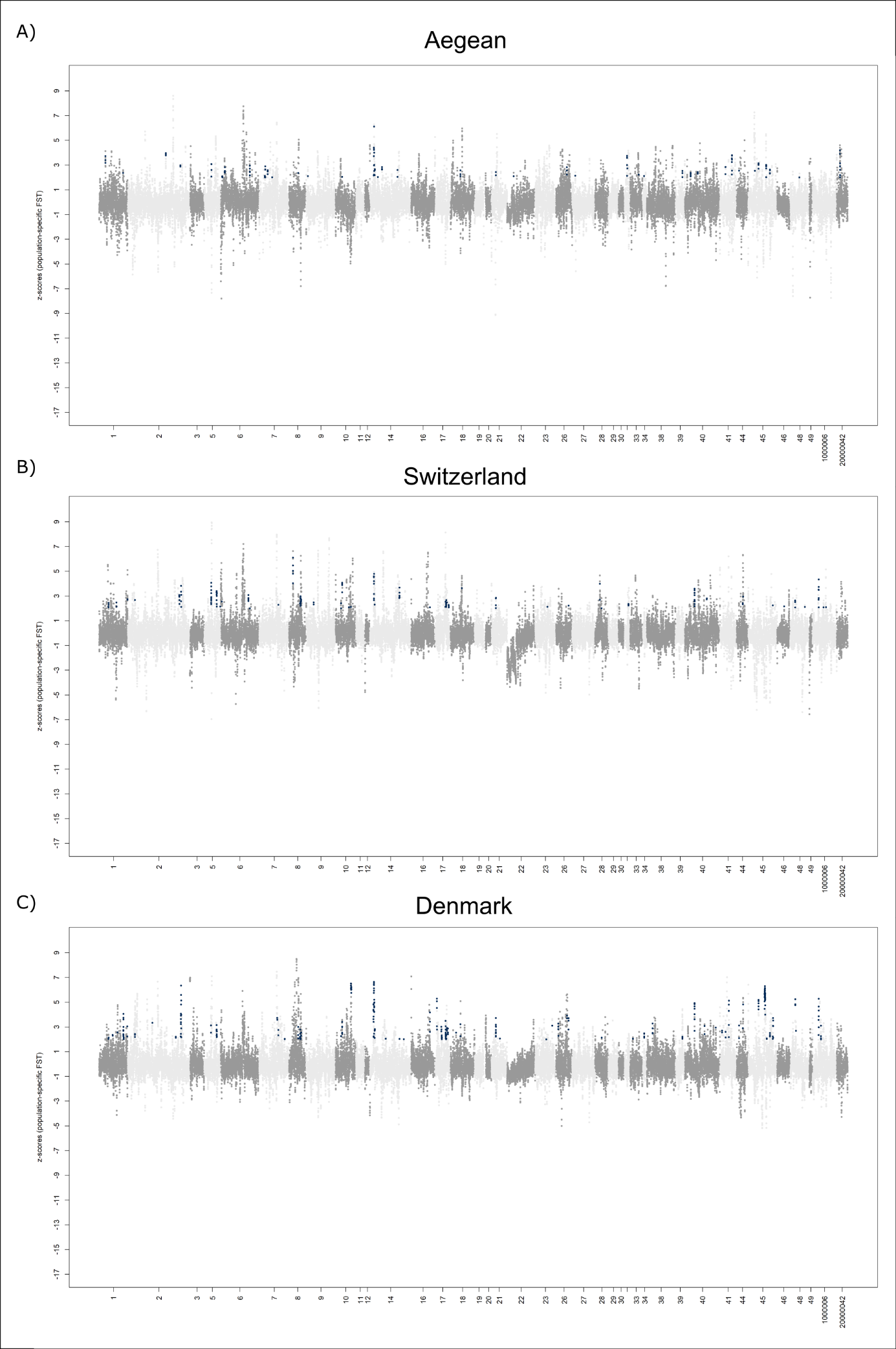

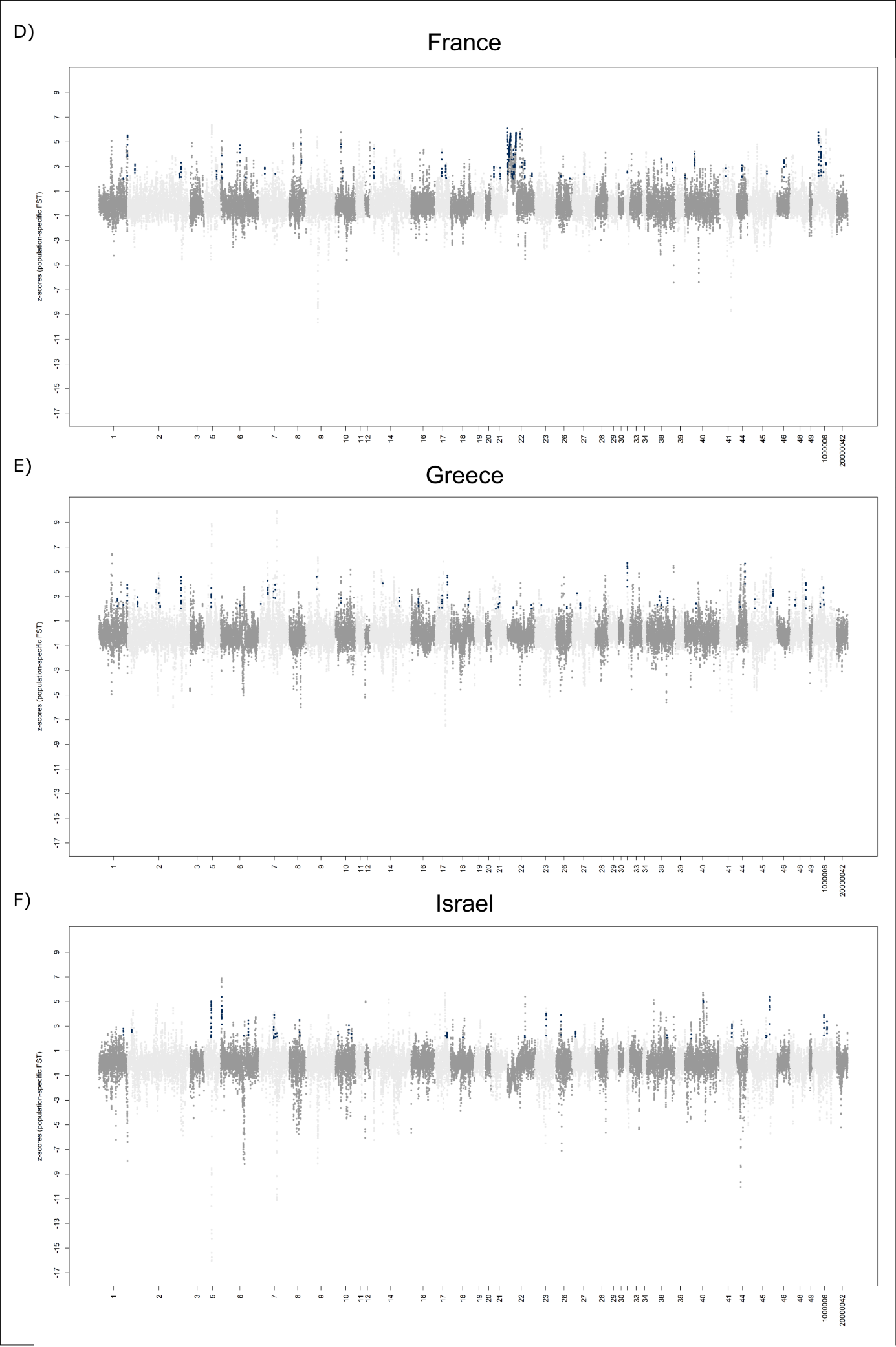

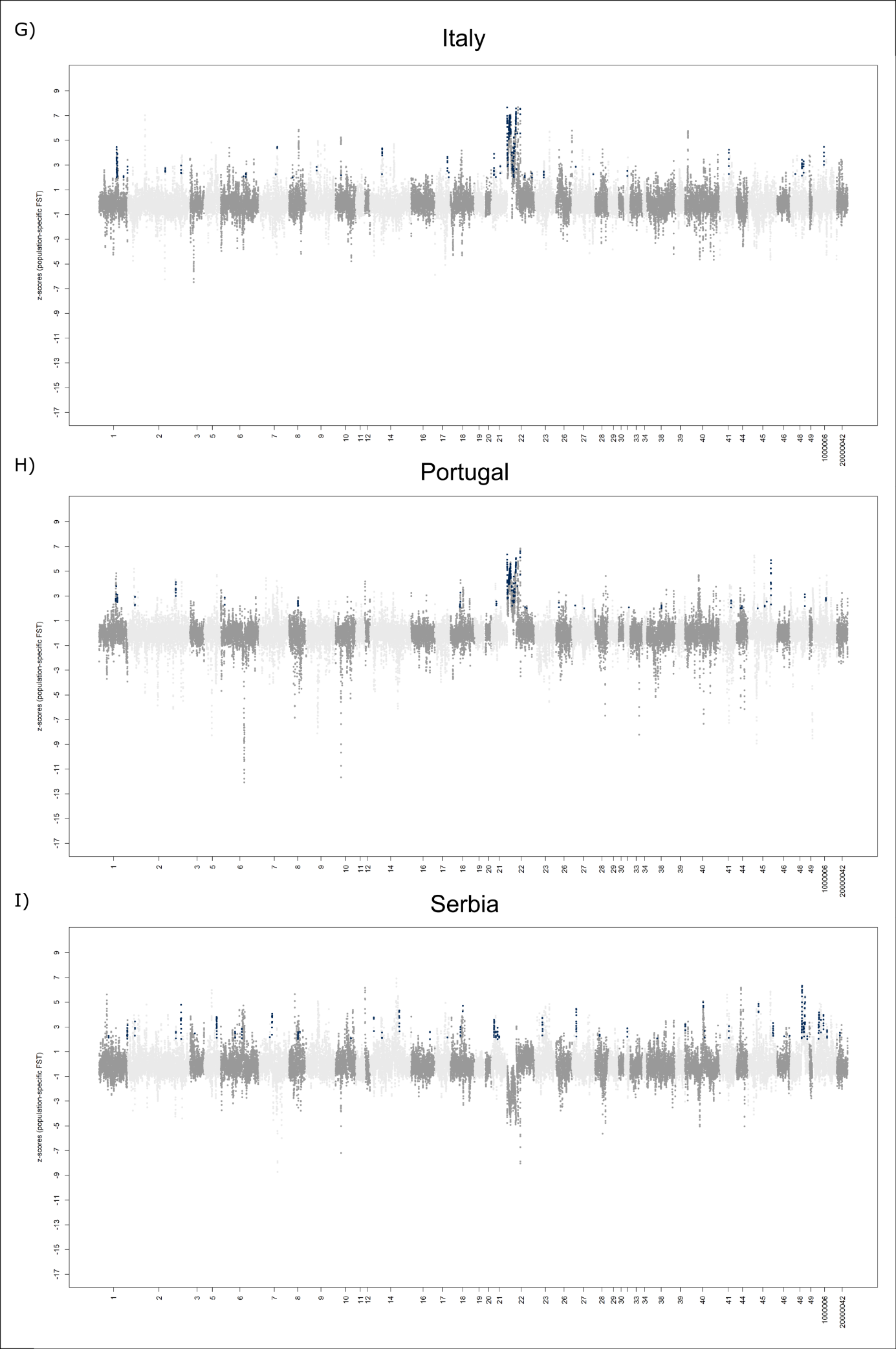

Figure S8 – Population-specific FST genome scans for each of the 9 populations. Each dot depicts an overlapping window of 100 kbp (step of 20 kbp. Highlighted in dark blue are the final outlier windows, i.e., overlap between outliers of population-specific FST and the ones of WZA. A switch between light and dark grey represents a change in the scaffold. The names of all the scaffolds are displayed on the x-axis.

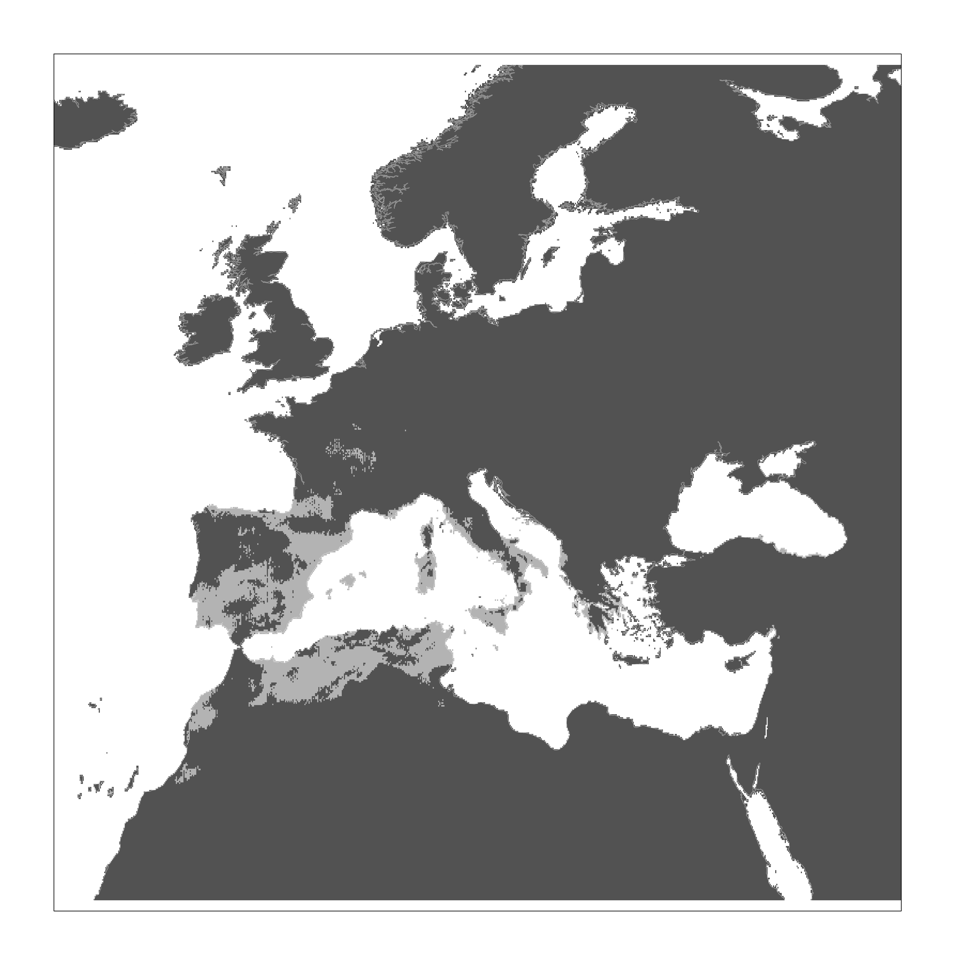

Figure S9 – Regions that were classified as suitable both during the LGM and today. Colours as above.

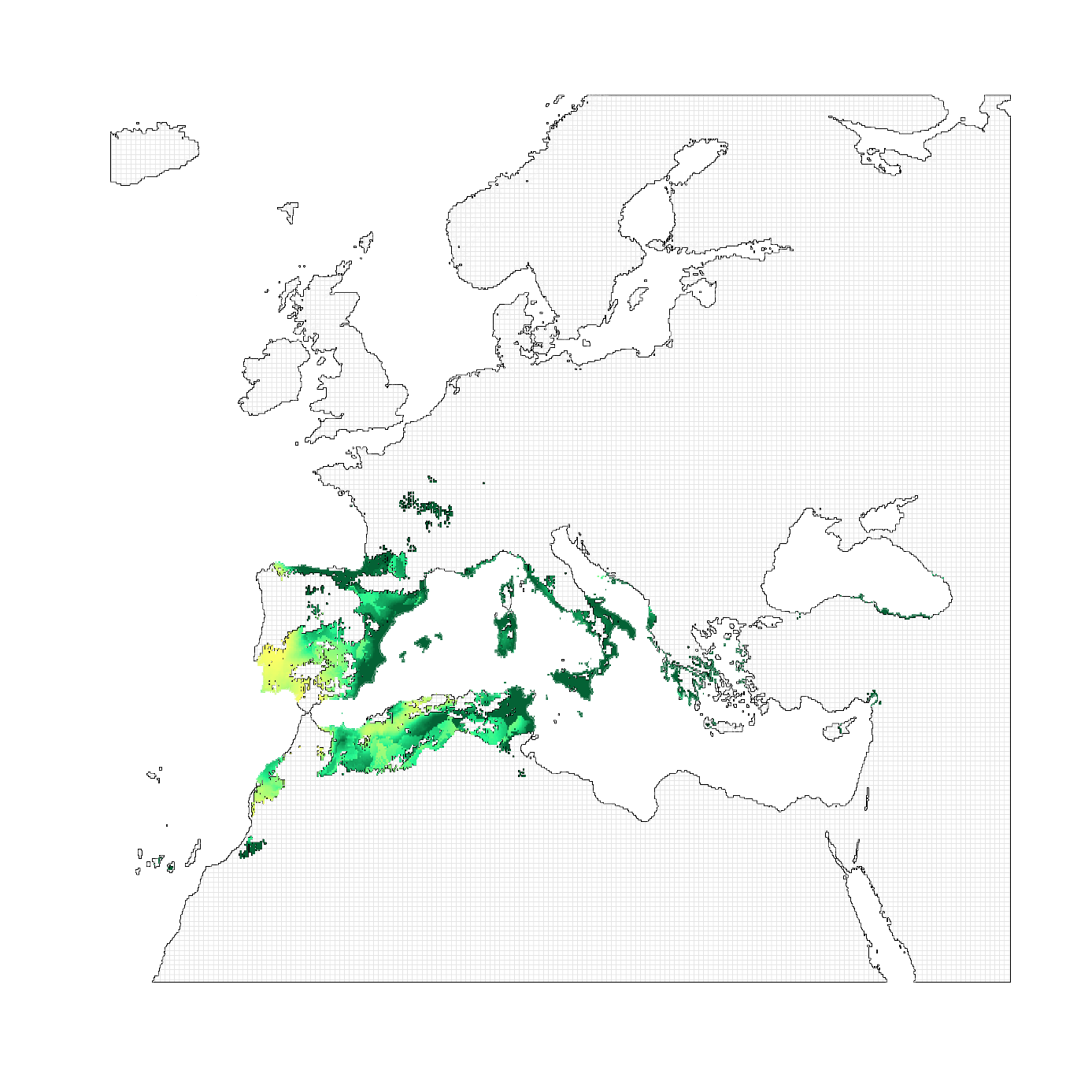

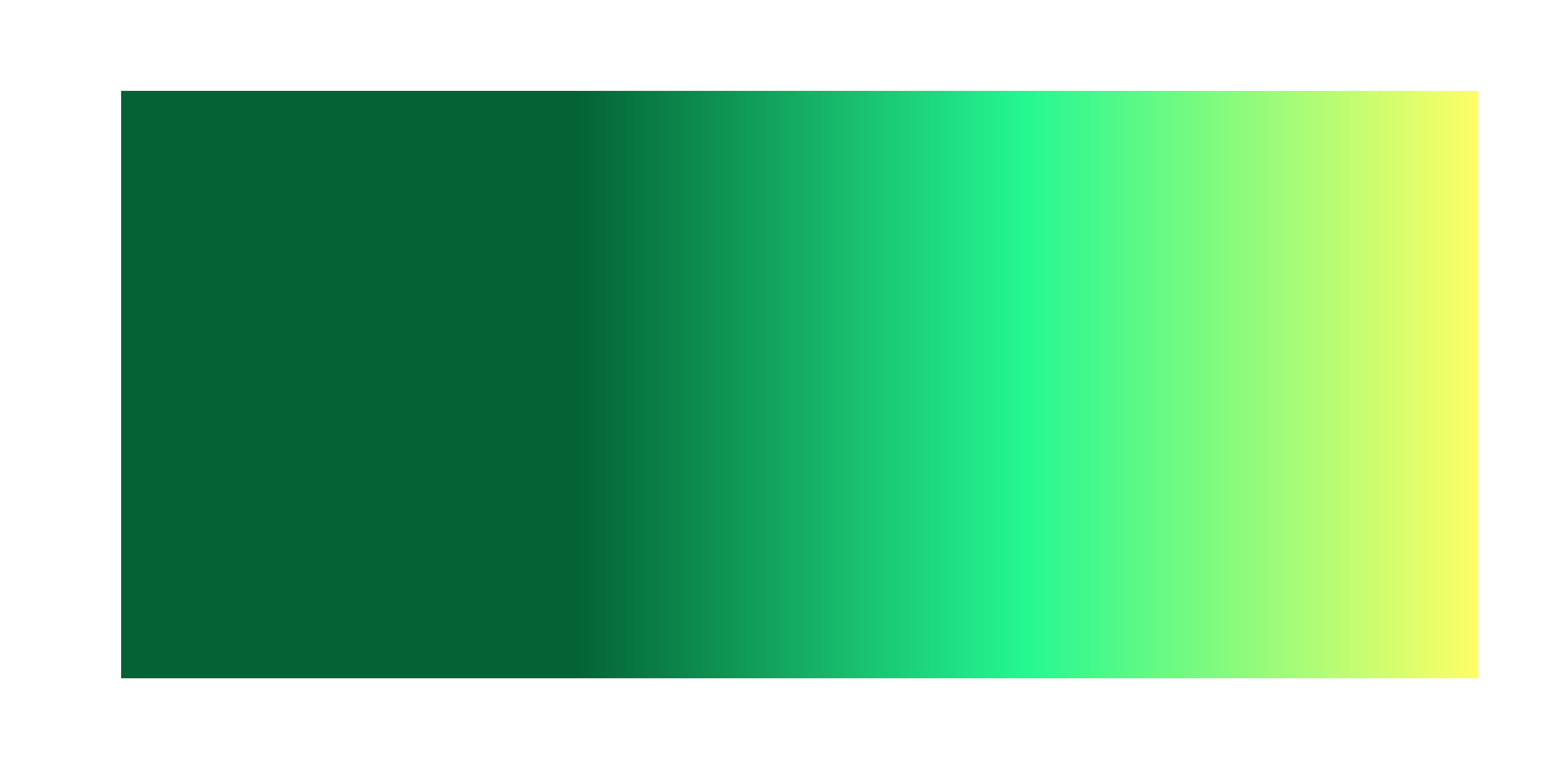

**0**

**1**

Figure S10 - Climatic shift among suitable regions both during the LGM and today. Map showing the shift of climatic conditions between the LGM and nowadays in the cells suitable at both time points. The light grey grid pattern shows unsuitable areas according to our SDM, while the rest is considered as suitable. Dark green to yellow gradient corresponds to the relative shift in conditions between the two time-points, with the highest changes in yellow.

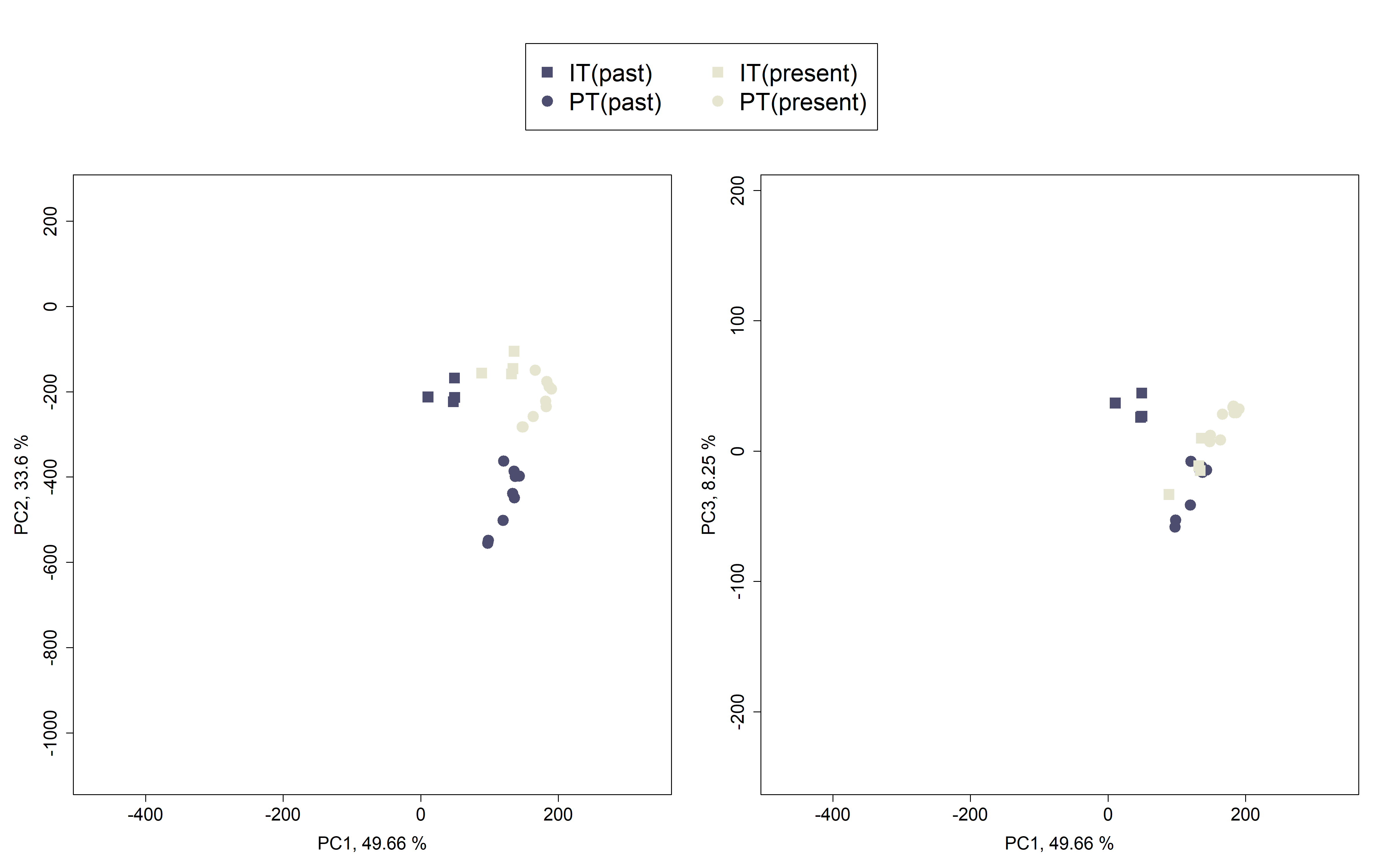

Figure S11 – Position in the 3D-climatic space of the climatic conditions during the LGM (in purple) and today (in light grey) of the Italian (squares) and Portuguese (round) sampling localities. We extract at the GPS coordinates the corresponding climatic conditions in the past and the present and projected them into the PCA made on bioclimatic variables from the entire study area (See Estimation of the shift of climate within suitable areas for details).

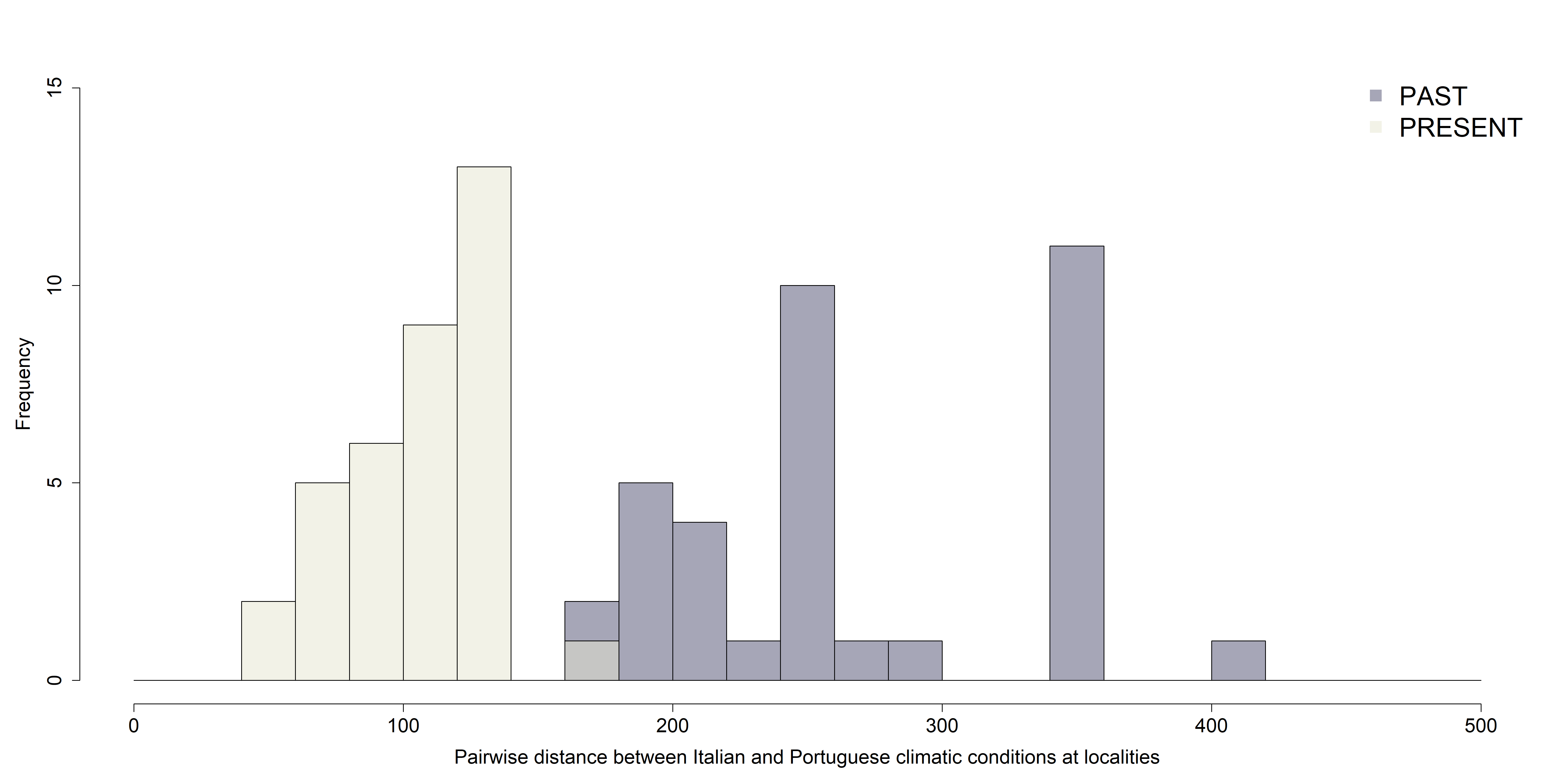

Figure S12 – Distribution of the Euclidean distances between all sampling localities of the two peninsulas during the LGM (purple) and the present time (light grey). This way, we quantified the climatic difference between the Italian and Portuguese sampling localities at the two different time points.

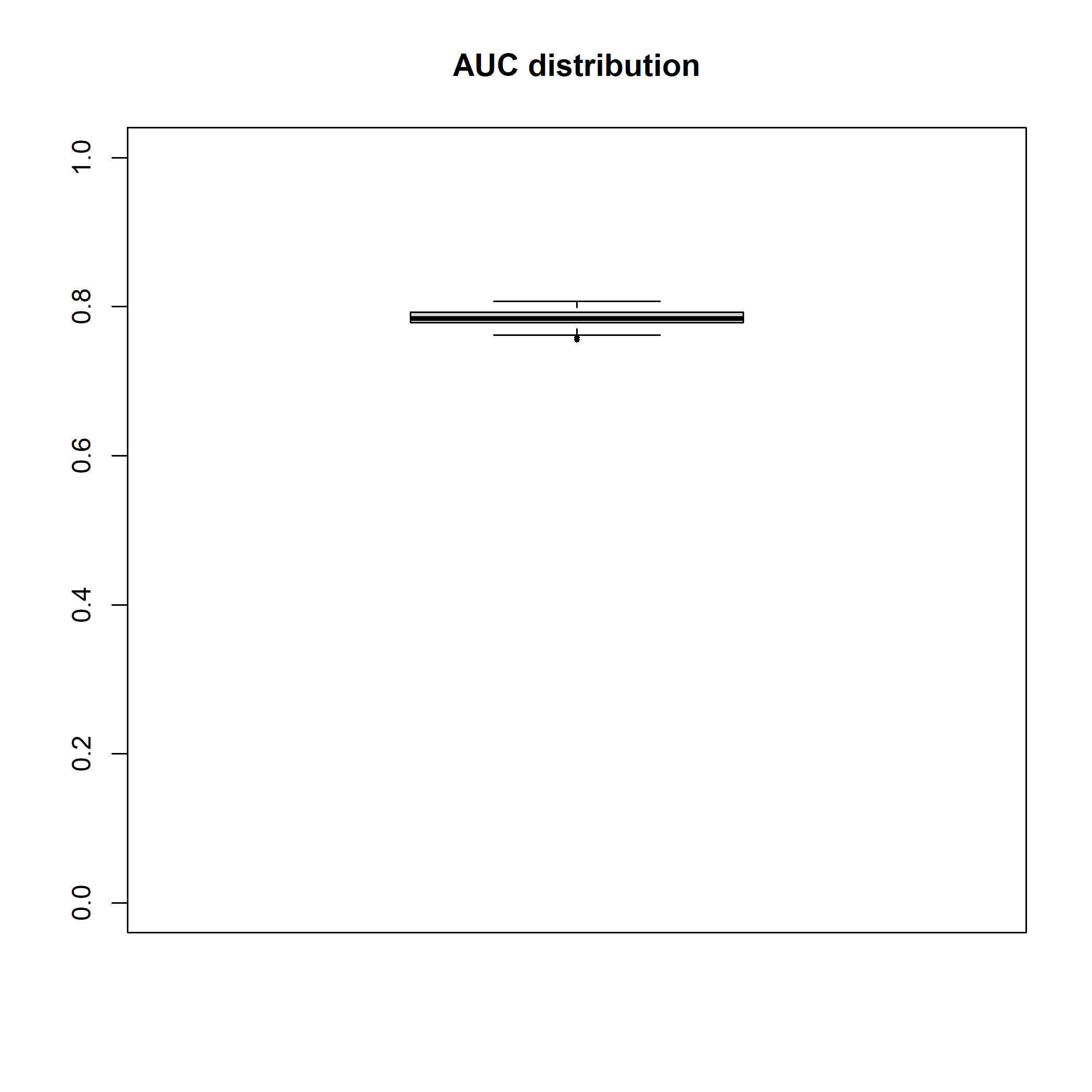

Figure S13 – Distribution of the AUC of the receiver operating characteristic (ROC) from our 100 MaxEnt models. The median was equal to 0.7842.

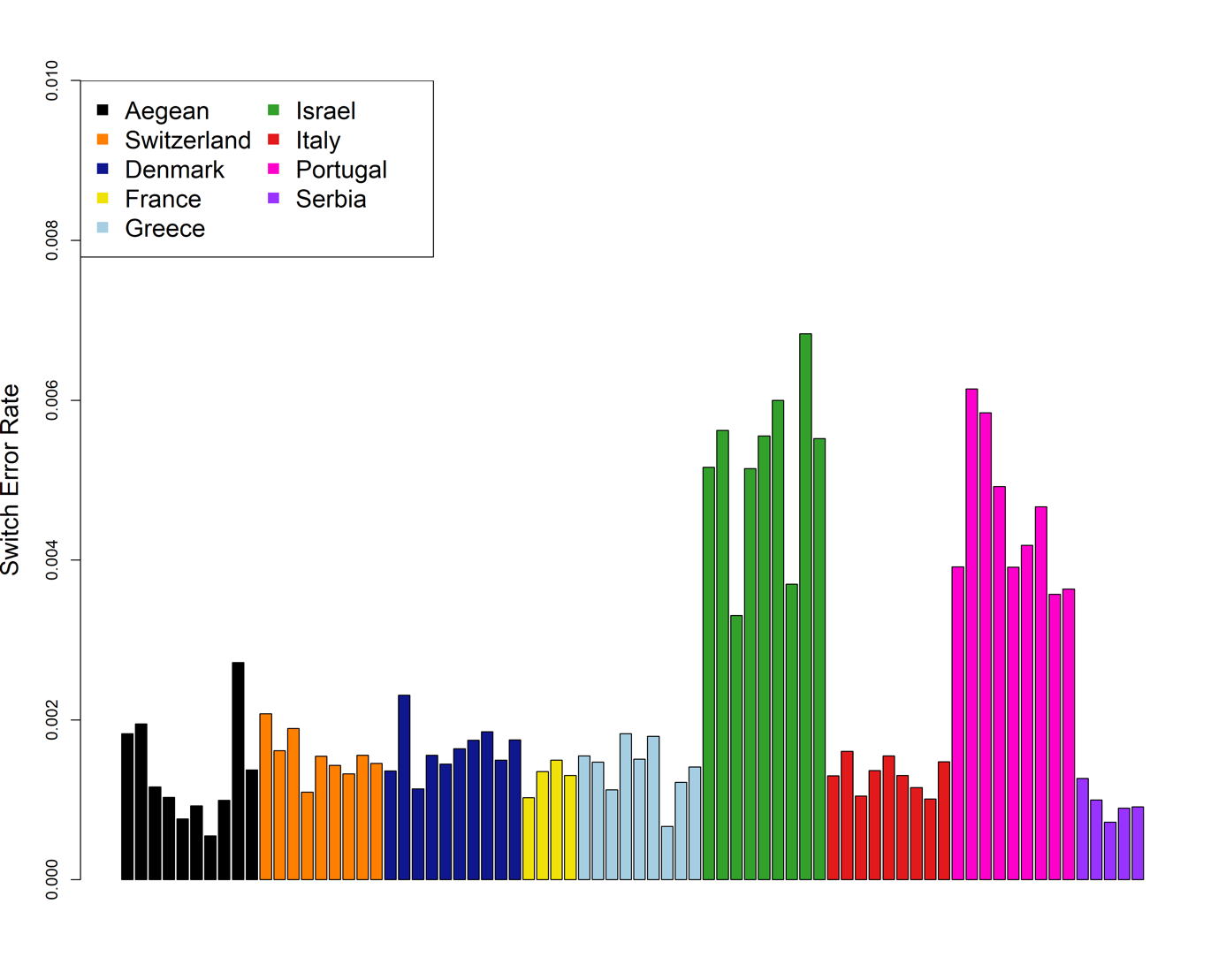

Figure S14 – Quantification of the Switch Error Rate resulting from the statistical phasing. We ran a statistical phasing of each individual without considering the read-based phasing from WhatsHap. This phasing was then compared to the “true” local phasing, inferred from the read-based approach (WhatsHap). The switch error rate between both phasing sets was estimated using the switchError code. Each bar represents one individual, coloured by population.

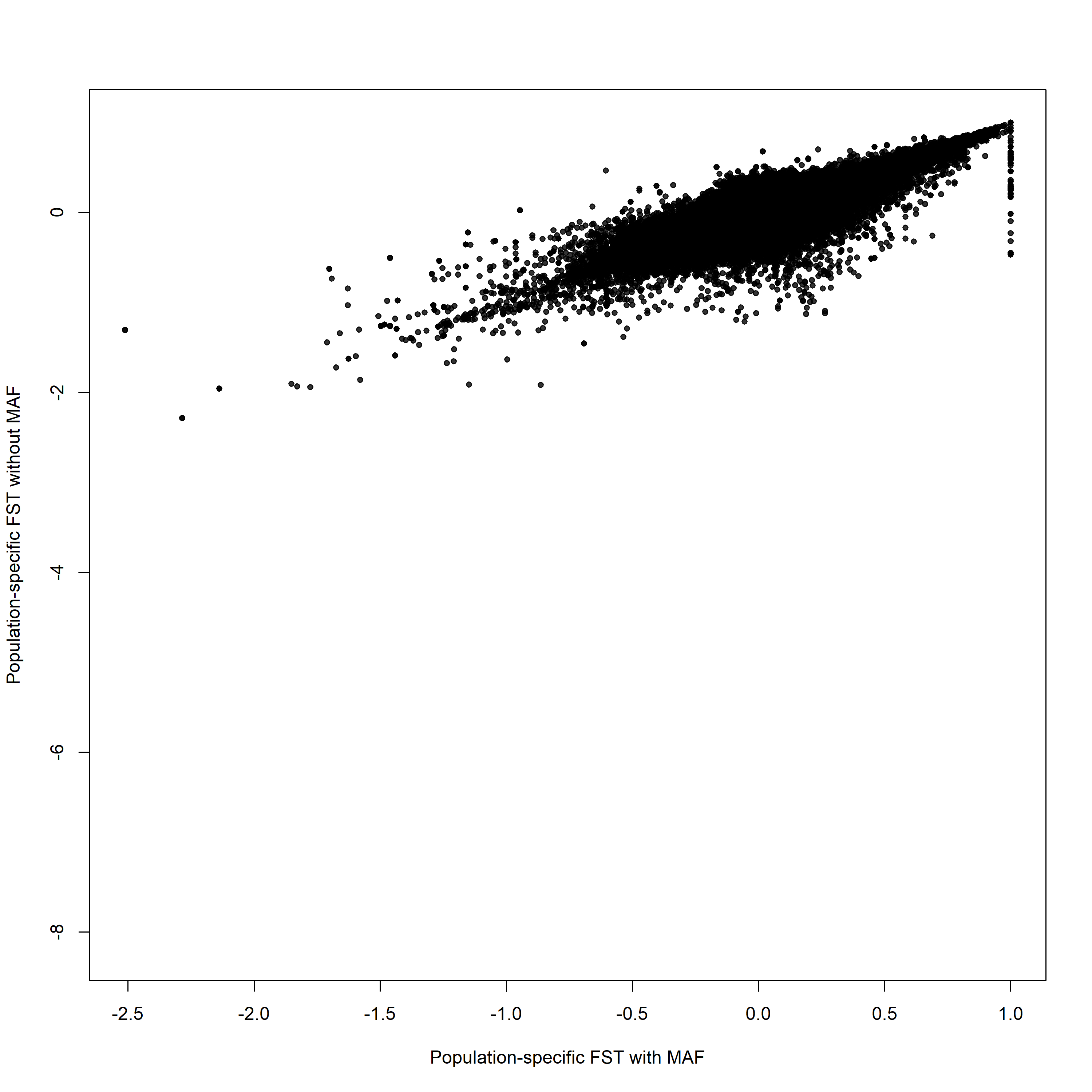
Figure S15 – Relationship between population-specific FST computed on SNP non-filtered for MAF and the one computed on SNP filtered on MAF. The first dataset was made of 12,309,943 SNPs; the second one of 4’689’284 SNPs. The correlation between the two statistics is 0.88.

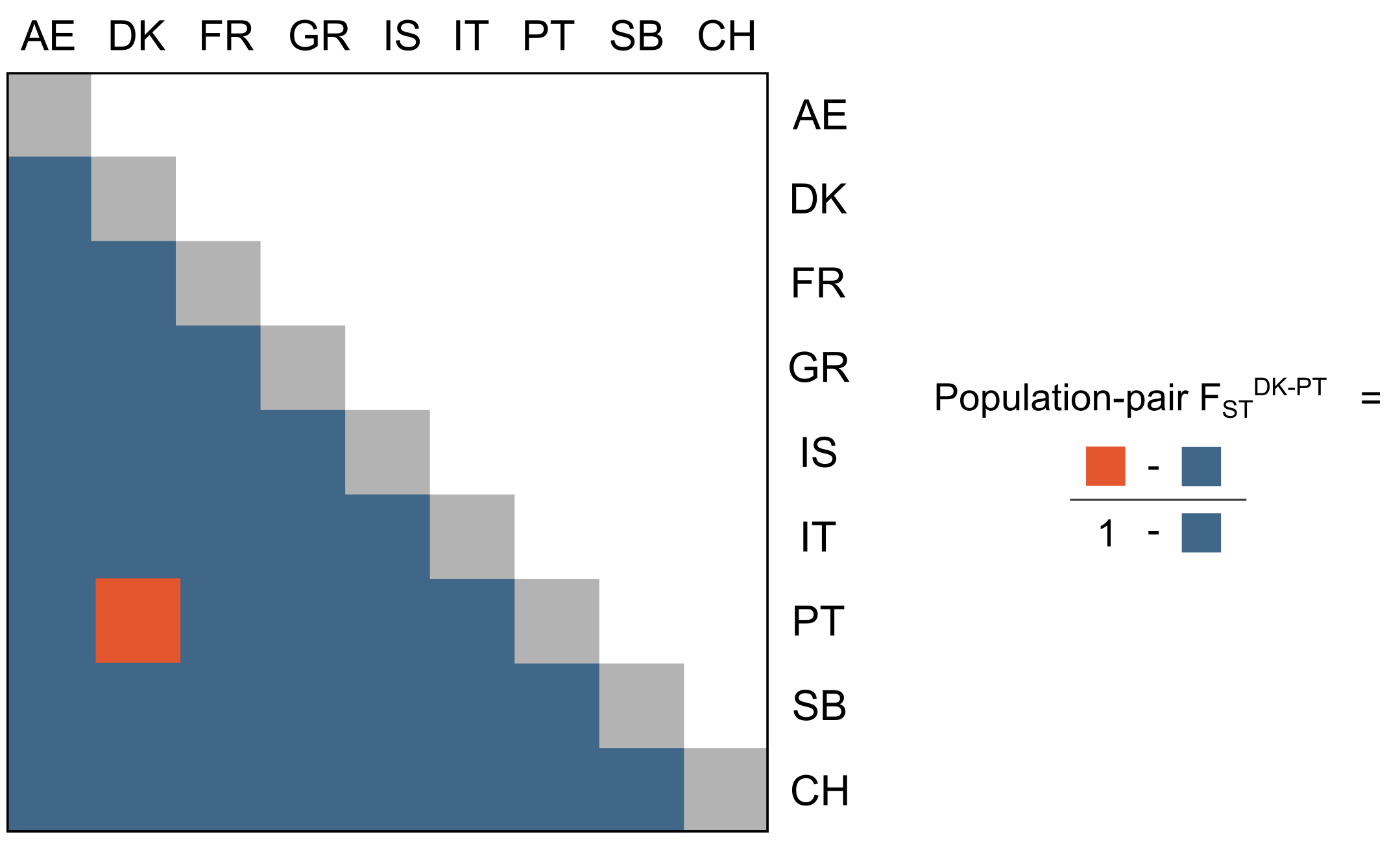

Figure S16 – Graphical representation of the calculation of population-pair FST between Denmark (DK) and Portugal (PT) populations. The matrix displays the mean allele sharing between all population pairs, with dimensions *n* × *n*, where *n* is the number of populations analysed.

### Supplementary Tables

Table S1 – Description of the samples used in this study. See supplementary material.

| **Population** | **Population-specific FST outliers** | | **Final outliers** | |
| --- | --- | --- | --- | --- |
|  | **Total number** | **Private** | **Total number** | **Private** |
| Aegean | 1673 | 883 | 126 | 67 |
| Switzerland | 1765 | 749 | 148 | 41 |
| Denmark | 2068 | 987 | 303 | 147 |
| France | 2087 | 769 | 348 | 72 |
| Greece | 1796 | 900 | 170 | 85 |
| Israel | 989 | 630 | 136 | 83 |
| Italy | 1824 | 629 | 295 | 66 |
| Portugal | 1521 | 521 | 243 | 54 |
| Serbia | 2084 | 966 | 265 | 129 |

Table S2 – Distribution of the different types of outlier windows among the nine populations. “Genome-scans outliers” windows describe the number of significant windows from population-specific FST (see Material and Methods section for details). “Final outliers” windows were obtained by retaining the overlap between the outlier from population-specific FST and the GEA approach (WZA). “Total number” refers to the number of windows detected in each population. “Private” represents the number of windows exclusive to each of the population.

| **Population** | | AE | CH | DK | FR | GR | IS | IT | PT | SB |
| --- | --- | --- | --- | --- | --- | --- | --- | --- | --- | --- |
| **Extracted genes** | In total | 70 | 91 | 133 | 147 | 104 | 64 | 120 | 102 | 102 |
|  | Private | 36 | 31 | 64 | 21 | 60 | 26 | 18 | 20 | 48 |

Table S3 – Number of genes in the windows putatively under selection in each population. Outlier regions were identified based on the concordant signal between the population-specific FST and the Weighted-Z Analysis. Genes that partially or fully overlapped the final set of outlier windows were considered as potentially involved in the local adaptation and therefore extracted. The extraction was based on the annotation of the barn owl genome from the NCBI (GenBank assembly accession: GCA_018691265.1 – RefSeq assembly accession: GCF_018691265.1).

|  | **Df** | **Variance** | **F** | **Pr(>F)** |
| --- | --- | --- | --- | --- |
| **Model** | 7 | 200214 | 1.5366 | 0.001 |
| **Residual** | 66 | 1228481 |  |  |

Table S7 – Results of the permutation test to assess the significance of the relationship between genotype and environmental predictors. A test statistic (F-statistic) was computed from the regression on the observed data. Afterwards, additional regressions were carried out 999 times on permuted rows of the response data (i.e., the genotype matrix), allowing the establishment of the empirical null distribution of the statistics, to which the observed statistic was compared (Borcard et al., 2011).

|  | **Df** | **Variance** | **F** | **Pr(>F)** | **Cumulative constrained variance [%]** |
| --- | --- | --- | --- | --- | --- |
| **RDA1** | 1 | 53639 | 2.8818 | 0.009901 | 26.79 |
| **RDA2** | 1 | 33597 | 1.8050 | 0.009901 | 43.57 |
| **RDA3** | 1 | 29862 | 1.6044 | 0. 009901 | 58.49 |
| **RDA4** | 1 | 22707 | 1.2199 | 0. 009901 | 69.83 |
| **RDA5** | 1 | 21184 | 1.1381 | 0.019802 | 80.41 |
| **RDA6** | 1 | 20130 | 1.0815 | 0.138614 | - |

Table S8 – Results of the permutation test to assess the significance of the individual axes from the Redundancy Analysis. To select the number of RDA axes we retained, we performed a permutation test for each axis (n = 100) by following the procedure Borcard et al. (2011) gave and implemented in the vegan R package through the anova.cca() function (Oksanen et al., 2020).
